# Unusual dopamine-mediated regulation of the phototransduction in lamprey compared to jawed vertebrates

**DOI:** 10.1101/2025.06.26.661820

**Authors:** D.A. Nikolaeva, A.Yu. Rotov, I. Yu. Morina, M.L. Firsov, I.V. Romanova, L.A. Astakhova

## Abstract

The vertebrate retina uses neurotransmitters to regulate its various functions and adjust vision according to the day/night cycle. Dopamine is probably one of the most important of these neurotransmitters. It is released by dopaminergic amacrine cells in the retina and exerts its regulatory effects, in part, through the cAMP pathway. It has been demonstrated that dopamine affects the phototransduction cascade in isolated amphibian rods. Furthermore, elevated intracellular levels of cAMP increase the light sensitivity of vertebrate rods and modulate the response of vertebrate cones. These effects can be triggered by dopamine receptors and adjust vision to daily light variations. Therefore, the evolution of dopamine loops in the retina is of interest, and the lamprey, being the most primitive vertebrate, could be valuable in this regard. In the present study, we examined whether the photoresponse properties of long (cone-like) and short (rod-like) photoreceptors in the river lamprey could be regulated by dopamine or cAMP level modulation. Using suction pipette recording, we demonstrated that dopamine slightly increased short photoreceptors sensitivity and it slowed the rising and falling phases of photoresponses in long photoreceptors and increased the integration time, with no effect on the sensitivity to brief flashes. The second part of our study — an immunohistochemical analysis of the lamprey retina — revealed that both D1 and D2 dopamine receptors are expressed in both types of lamprey photoreceptors. Our results suggest that the regulation of photoreceptor functions by the neurotransmitter dopamine originated in the early stages of vertebrate evolution, specifically during the Cambrian period.

**Summary:** The lamprey, a primitive vertebrate, is a valuable object for studying the evolution of dopamine loops in the vertebrate retina. This study shows that photoresponse properties of lamprey photoreceptors are regulated by dopamine in a different way compared to gnathostomes.

## Introduction

The vertebrate retina adapts its functions to the day/night cycle in order to regulate vision. Neurotransmitters are vital for this process, with dopamine being one of the most significant (Witkovsky, 2004; Roy and Field, 2019). Dopamine is released in the retina by a specialised subset of cells known as dopaminergic amacrine cells in response to light (Dacey, 1990), a process controlled by photosensitive retinal cells, primarily rods (Pérez-Fernández et al., 2019). The level of dopamine in the retina fluctuates cyclically, increasing during the day and decreasing at night. Dopamine in the vertebrate retina regulates gap junctions between different types of retinal cells, forming both homotypic and heterotypic coupling (Ribelayga et al., 2008; Xin and Bloomfield, 1999; Hu et al., 2010; Mills et al., 2007), and the transmission of signals through glutamate and GABA receptors in retinal neurons (Bergum et al., 2024; Maguire and Werblin, 1994). It also acts on numerous other cellular targets (reviewed in Popova, 2020; Goel and Mangel, 2021). With regard to photoreceptor cells, it has been demonstrated that dopamine is involved in signal transduction between photoreceptors via gap junctions (Krizaj et al., 1998; Ribelayga et al., 2008; Jin et al., 2015); the regulation of synaptic transmission between rods, cones, and horizontal cells (Stella and Thoreson, 2000; Thoreson et al., 2002); hyperpolarisation-activated currents in rod photoreceptors (Akopian and Witkovsky, 1996; Kawai et al., 2011); retinomotor movements (Dearry and Burnside, 1986; Dearry et al., 1990); and the phagocytosis of outer segment membrane discs (Reme et al., 1986; Besharse et al., 1979). We have previously demonstrated that dopamine and dopamine receptor agonists affect the functioning of the phototransduction cascade in isolated amphibian rods, modulating their sensitivity and response kinetics (Nikolaeva et al., 2018). Dopamine decreases rod sensitivity by reducing the activation rate of the cascade and, to a lesser extent, by speeding up its deactivation. Furthermore, as previously demonstrated, elevated intracellular levels of cAMP significantly increase the light sensitivity of amphibian rods (Astakhova et al., 2012) and markedly modulate the response and dark current recovery in fish cones (Sitnikova et al., 2020; Chrispell et al., 2022). These regulatory loops could serve as the basis for signalling from dopamine receptors when adapting the phototransduction cascade to diurnal variation in ambient light.

Extracellular dopamine acts on retinal neurons through specific receptor proteins. Based on their ability to modulate cAMP production, dopamine receptors can be divided into two classes: D1-type and D2-type (Spano et al., 1978; Kebabian and Calne, 1979; Gingrich and Caron, 1993). Activation of these receptors triggers a variety of intracellular events, which may or may not be related to changes in cAMP concentration within a cell. Most major retinal cell types appear to express dopamine receptors. D1-type receptors have been identified in horizontal, bipolar, ganglion, amacrine, and pigment epithelial cells (Versaux-Botteri et al., 1997; Veruki and Wassle, 1996; Mora-Ferrer et al., 1999; Farshi et al., 2016). In the retinas of vertebrates, D2-type receptors are expressed by both classes of photoreceptor (Dearry et al., 1990; Muresan and Besharse, 1993; Wagner et al., 1993), as well as by horizontal cells (Yazulla and Lin, 1995) and by bipolar, ganglion and amacrine cells (Veruki et al., 1997; Wagner et al., 1993). All of this knowledge about the functions of dopamine, its signalling and the distribution of dopamine receptors in the retina has been obtained from different classes of vertebrates whose retinas and cell types are contemporary from an evolutionary standpoint. However, it would be interesting to shed light on the evolution of dopamine loops in the retina. In this context, the lamprey’s retina could be a useful model. Lampreys belong to the jawless class of vertebrates and are the most primitive currently existing vertebrates; their ancestors diverged from other vertebrate species over 500 million years ago (Bayramov et al., 2018; Kuraku and Kuratani, 2006). Nevertheless, lampreys have well-designed, camera-type eyes with an orderly, layered retina and neuron types that are conserved between jawless and jawed lineages. These include photoreceptors, bipolar, horizontal, amacrine and ganglion cells (Fain, 2019; Wang et al., 2024). The lamprey retina is duplex and has distinct rod- and cone-like photoreceptors (Asteriti et al., 2015; Govardovskii et al., 2020). Nevertheless, the retinas of jawless and higher vertebrates exhibit notable differences. In particular, lamprey rods are morphologically similar to cones; the outer segments of both consist of invaginating lamellae formed by the plasma membrane. Additionally, retinal ganglion cells are located on both sides of the inner plexiform layer.

The present study investigated whether the photoresponse properties of the long (cone-like) and short (rod-like) photoreceptors in the river lamprey retina could be regulated by dopamine and changes in intracellular cAMP levels. The second part of the study focused on the immunohistochemical evaluation of dopamine receptor expression in the various layers and cell types of the lamprey retina. This remains unclear to our knowledge.

## Materials and methods

### Experimental animals and dissection procedures

River lampreys (*Lampetra fluviatilis*) were caught in the rivers of the St. Petersburg vicinity in November 2020, 2021 and 2022, and were taken to the vivarium at the Sechenov Institute of Evolutionary Physiology and Biochemistry (IEPhB). They were kept in a cold room at 4–6 °C in large basins containing filtered, well-aerated water (25–30 litres per animal) in the dark. Adult marsh frogs (*Pelophylax ridibundus*) were caught in the wild in southern Russia (the Astrakhan region) in September 2022 and 2023. They were then transported to the IEPhB vivarium, where they were kept under a layer of water in large tanks in refrigerators (at 4–6 °C) in the dark. The frogs were kept without food because their metabolic rate was greatly reduced in these conditions. Two days prior to the experiment, the frogs were removed from the refrigerator and placed on a natural day/night cycle. Prussian carp (*Carassius gibelio*) were obtained from local hatcheries and kept in a 40 L aerated aquarium containing 5–7 fish at a time. The aquarium was kept in a laboratory room at 21–23 °C and maintained under a 12-hour on/12-hour off lighting cycle. The fish were fed spirulina-based dry fish food.

For the electrophysiological studies and tissue extraction for Western blotting, all animals were dark-adapted overnight prior to each experiment. For histology, after overnight dark adaptation, the animals were exposed to moderate white illumination for 30 minutes to ensure tighter contact between the retina and the retinal pigment epithelium (RPE) and reduce detachment during dissection and fixed tissue processing. They were then euthanised by decapitation and destruction of the spinal cord. The eyes were enucleated under dim red light. All subsequent dissection procedures (e.g. cutting the eyeball, removing the lens and cornea, and extracting the retina from the eyecup) were performed using a binocular magnifier with infrared illumination. The retinal samples were then used for either photocurrent recording experiments or immunohistostaining (see below for details of the methods).

The handling of experimental animals complied with the requirements of European Directive 2010/63/EU and the recommendations of the Bioethics Committee of the Sechenov Institute of Evolutionary Physiology and Biochemistry of the Russian Academy of Sciences (Permit #8/2021, issued on 26 August 2021).

### Electrophysiology

#### Preparation and solutions

Electrophysiological experiments were conducted on the retinas of lampreys and fish. A piece of the isolated retina was transferred into a drop of Ringer’s solution (see composition below), shredded into small fragments with fine forceps and gently pipetted several times to obtain isolated photoreceptors. As the lamprey vitreous humour is quite viscous, the retina was exposed to 0.25 mg/ml hyaluronidase for 2 minutes prior to shredding, to prevent clogging of the glass micropipettes. The resulting suspension of small retinal pieces and isolated photoreceptors was then transferred to a perfusion chamber in the experimental setup.

The retinal cells in the recording chamber were perfused constantly with the same Ringer’s solution that was used for dissection and storage of the retinal sample. The composition of the Ringer’s solution was species-specific. Lamprey: NaCl 120, KCl 3.6, MgSO₄·7H₂O 1.2, CaCl₂ 1.1, NaHCO₃ 22.6, HEPES 10, glucose 6; pH adjusted to 7.6. Frog: NaCl 90, KCl 2.5, MgCl₂ 1.4, CaCl₂ 1.05, NaHCO₃ 5, HEPES 5, glucose 10 and EDTA 0.05, with the pH adjusted to 7.6. Fish’s Ringer’s solution contained the following in mM: NaCl 102, KCl 2.6, MgCl₂ 1, CaCl₂ 1, NaHCO₃ 28, HEPES 5 and glucose 5, with the pH adjusted to 7.8.

The drug-containing solutions used in the experiments were 10 µM forskolin and 50 µM and 250 µM dopamine, dissolved in Ringer’s solution. The forskolin solution was prepared from a fresh stock solution (10 mM) in dimethyl sulfoxide (DMSO) on the day of the experiment. The final concentration of DMSO in the perfusion solution was less than 0.1%, which had no negative effect on photoreceptor viability or electrical activity. The dopamine solution was made up from powder immediately prior to application to the tested photoreceptor, since dissolved dopamine is susceptible to oxidation over time. All chemicals were purchased from Sigma-Aldrich (St. Louis, MO, USA).

The preparation procedures and subsequent electrophysiological recordings were carried out at a room temperature of 17–19 °C.

#### Single-cell recordings and experimental protocol

The responses of single photoreceptors were studied using the suction micropipette technique (Baylor et al., 1979). Details of our suction rig and how it works are given in our previous publications (Astakhova et al., 2008, 2015).

Glass suction micropipettes were pulled using a micropipette puller (P-97 Flaming/Brown Micropipette Puller, Sutter Instrument, USA). In a perfusion chamber, a photoreceptor was pulled into the micropipette by its outer segments. We usually recorded from single photoreceptors protruding from the edge of the lamprey retina, or from isolated cones (green-sensitive members of double cones) in the case of the fish retina.

The cells were stimulated with 10 ms (rods) or 2 ms (cones) of LED light at λ_max_ = 525 nm. Stimulation using red (λ_max_ = 630 nm), green (λ_max_ = 525 nm) or blue (λ_max_ = 460 nm) LEDs in the second channel allowed us to distinguish the green-sensitive fish cones unambiguously from the red- and blue-sensitive ones. The emission spectra of all the light stimuli used in the setup were recorded using a USB4000 spectrometer (Ocean Optics, USA). LED intensities were measured at the same positions where the preparations were located using an OPT-301 optosensor (Burr-Brown Corporation, USA). Flash intensity was regulated using switchable neutral density filters inserted into the beam. Photoresponses were recorded at 100 Hz for rods and 500 Hz for cones, and were low-pass filtered at 30 Hz using an eight-pole analogue Bessel filter. Data acquisition, stimulus timing and flash intensity were controlled by LabVIEW hardware and software (National Instruments, Austin, TX).

A photoreceptor placed in the micropipette was stimulated by light stimuli, which elicited saturated and fractional responses (approximately one-quarter and one-half saturated). This set of responses was recorded in Ringer’s solution. Then, the perfusion solution was replaced with a solution containing forskolin or dopamine (the time to completely change the solution in the sample holder is approximately 2.5 to 3 minutes). The responses were recorded according to the light stimulation protocol after 20 and 30 minutes of incubation. The same experimental protocol was used for control experiments conducted without the addition of drugs, to assess the extent to which the metabolic rundown of cells affected response amplitude over time.

#### Data processing and statistical analysis

Data captured in the setup were processed using custom software written in LabVIEW. The initial and falling phases of photoresponses were fitted using custom-made Python software. All light response parameters in control and drug-containing solutions were self-normalized using the values achieved from the same cell during pre-incubation in pure Ringer solution.

The normality of datasets from lamprey and Carassius photoreceptor light responses was confirmed using the Shapiro–Wilk test. Parameters in the control and drug-containing solutions were compared using either a one-way ANOVA with a post hoc Dunnett’s multiple comparisons test or an unpaired t-test. A p-value of less than 0.05 was taken to indicate statistical significance.

#### SDS-polyacrylamide gel electrophoresis (SDS-PAGE), immunoblotting

The dissected retinas were homogenised in the ratio 1:10 in the lysis buffer containing the following: 20 mM Tris-HCl (pH 7.5), 150 mM NaCl, 2 mM EDTA, 2 mM EGTA, 0.25 M sucrose, 0.5% Triton X-100, 0.5% sodium deoxycholate, 15 mM NaF, 10 mM sodium glycerophosphate, 10 mM sodium pyrophosphate, 1 mM Na3VO4, 1 mM phenylmethylsulfonyl fluoride (PMSF), 0.02% NaN3, and the protease inhibitor cocktail (Sigma-Aldrich, USA). The homogenates were then centrifuged for 15 minutes at 11,000 g and 4 °C. The resulting supernatant was collected, and the proteins in the sample were solubilized by boiling in Laemmli sample buffer. The SDS-PAGE and immunoblotting protocols have been described previously (Derkach et al., 2019). Briefly, the protein samples were fractionated by SDS-PAGE in 10% and 12% polyacrylamide gels, then transferred to 0.22 μm nitrocellulose membranes (GE Healthcare, Amersham Biosciences AB, UK) by electroblotting (300 mA for 1 hour) in a mini trans-blot module (“Bio-Rad Laboratories Inc.”, USA). Non-specific binding was blocked by incubating the membranes in 5% non-fat dry milk diluted in TBST (0.1% Tween-20 in Tris-buffered saline) for 1 h at room temperature. The membranes were then incubated overnight at 4 °C with primary antibodies diluted in blocking solution: rabbit anti-D1 dopamine receptor (D1RD; Abcam; 1:1000) or rabbit anti-D2(L/S) dopamine receptor (D2RD; Millipore; 1:500). After removing the primary antibodies, the membranes were incubated with horseradish peroxidase (HRP)-conjugated goat anti-rabbit IgG (Sigma; 1:10,000 dilution) for 1 h at room temperature. The membrane was washed with TBST three times for 10 minutes each time. The Novex ECL Chemiluminescent Substrate Reagent Kit (Invitrogen, USA) and premium X-ray film (GE Healthcare, UK) were used to visualize the bands.

#### Immunohistochemistry

The slides with retina sections were washed with PBS, treated with a 0.6% hydrogen peroxide solution in PBS for 30 minutes to block endogenous peroxidase activity, then washed with PBS for 15 minutes. Finally, they were washed with a PBS solution containing 0.1% Triton X-100 (PBST) for a further 30 minutes. The sections were then incubated for 1 hour in a blocking solution (a mixture of 3% goat serum and 2% bovine serum in PBST). The sections were then incubated with either the primary rabbit anti-D1 dopamine receptor antibody (Abcam; dilution 1:300) or the primary rabbit anti-D2 receptor antibody (Millipore; dilution 1:200) in a 2% blocking solution for 48 hours at 4 °C. After washing in PBST for 40 minutes, the sections were incubated for one hour in PBST with goat anti-rabbit biotin-conjugated IgG (Vector Laboratories, Inc.) at a dilution of 1:600. After washing in PBS, the sections were incubated for 1 hour in a streptavidin-peroxidase solution at a dilution of 1:1000 (Sigma, USA) in PBS. After washing in PBS, the sections were treated with a solution of 0.05% diaminobenzidine (Sigma-Aldrich, St. Louis, MO, USA) and 0.03% hydrogen peroxide in PBS. The reaction was stopped using distilled water. After washing, the sections were mounted in glycerol under coverslips. Some sections were stained with haematoxylin. The specificity of the immunohistochemical reaction was verified using a negative control (samples without primary or secondary antibodies; see Fig. S7).

#### Microscopy

Micrographs of retina sections were obtained using a Carl Zeiss Imager A1 microscope with transmitted light (Axio Vision 4.7.2 software, Carl Zeiss, Jena, Germany) and an Axiocam 712 video camera (using Zen 3.4 Blue Edition software, Carl Zeiss, Oberkochen, Germany).

The retina of vertebrates contains the five conventional classes of neurons (photoreceptors, horizontal cells, bipolar cells, amacrine cells and ganglion cells), which are organised into three main nuclear layers and two plexiform layers containing axons and dendrites. While the lamprey retina generally follows this organisational pattern, it has several specific morphological features (Fain, 2020). In gnathostomes, ganglion cell axons converge at the vitreal edge of the retina to form the optic nerve, which extends into the central nervous system. In lampreys, however, the ganglion cells’ axons form an optic fibre layer (OFL) located between the inner nuclear layer (INL) and the inner plexiform layer (IPL) (Holmberg, 1978). Most of the lamprey retina’s ganglion cells are displaced to the INL, with only a few observed within the IPL to form the inner ganglion cell layer (IGCL) (Fritzsch & Collin, 1990; Jones et al., 2009). Additionally, horizontal cells in the lamprey retina form two distinct sublayers (outer and inner horizontal cells, OHC and IHC), rather than being integrated into the INL as they are in gnathostomes (Öhman, 1976). Therefore, in the following text, we will use a different nomenclature for the layers of the lamprey retina compared to the retinas of fish and frogs.

#### Online supplemental material

Fig. S1 shows the results on 50 and 250 μM dopamine effects on the dark current, light sensitivity and how fast lamprey short photoreceptors react to light after about 20 minutes (1st time point in dopamine). Fig. S2 presents the results of experiments conducted on the impact of 50 and 250 μM dopamine on the dark current, light sensitivity and photoresponse kinetics of lamprey long photoreceptors following a 20-minute exposure period. Fig. S3 shows the effects of 10 μM forskolin on the dark current, light sensitivity and photoresponse kinetics of lamprey short photoreceptors after approximately 20 minutes’ exposure. Fig. S4 shows the effects of 10 μM forskolin on the dark current, light sensitivity and photoresponse kinetics of lamprey long photoreceptors after approximately 20 minutes’ exposure. Fig.S5 shows the effects of 250 μM dopamine on the dark current, light sensitivity and photoresponse kinetics of green-sensitive cones of Carassius after approximately 20 minutes exposure. Fig. S6 shows Western blotting results for lamprey and mouse retinas, and Fig. S8 shows Western blotting results for carassius, frog and mouse retinas. Fig. S9 shows the negative control for immunohistochemical reactions in the retinas of lampreys, carassius and frogs.

## Results

### Results

The aim of this study was to investigate the potential role of dopamine and the cAMP signalling pathway in the phototransduction cascade in two types of photoreceptor, long and short, in the retina of the river lamprey. Given the distinct physiological characteristics of these photoreceptors — long photoreceptors exhibit fast photoresponses and relatively low light sensitivity, resembling the cones of higher vertebrates, while short photoreceptors display slow photoresponses and high light sensitivity, becoming saturated at low light levels and resembling the rods of higher vertebrates — one might expect different contributions from the dopamine- and cAMP-mediated regulatory loops.

### Control experiments

In our experiments, we applied both dopamine and forskolin to different types of photoreceptor using a long-lasting protocol (30-minute perfusion or more) with a solution containing the tested pharmacological agents for each cell. To account for potential changes in the main photoresponse parameters over time, we performed control experiments in which each photoreceptor type was kept in Ringer’s solution for the same period. The photoresponses recorded in these control experiments at 20 and 30 minute time points were used to extract ‘control’ parameters for further statistical analysis.

### The effects of dopamine on the photoresponses of long and short photoreceptors in the river lamprey

#### Short photoreceptors

Electrophysiological experiments were carried out according to the established protocol. The first set of photoresponses was recorded in Ringer’s solution. After a 20- or 30-minute perfusion with a dopamine-containing solution (50 or 250 μM), the responses were recorded again using the same light stimulation parameters. The tested sets of responses included the following: 1) response to a saturating flash, 2) responses to a near half-saturating flash, and 3) responses to a near quarter-saturating flash. During data processing, the responses to non-saturating flashes were normalised to the dark current value at the corresponding time point. Then, the obtained values were compared for Ringer’s solution and for 20 and 30 minutes after dopamine application. The amplitudes of the normalised non-saturating responses were used to measure photoreceptor sensitivity, and the kinetic parameters of the responses were also compared. The magnitudes of saturated responses under different experimental conditions were compared to assess dark current.

Figure 1 shows a typical set of responses from lamprey short photoreceptors to saturating, nearly half-saturating, and quarter-saturating flashes, both in Ringer’s solution and 30 minutes after dopamine exposure. No significant impact on the dark current (i.e. the amplitude of saturated responses) or light sensitivity was observed in response to near quarter-saturating flashes. However, we observed a small but statistically significant increase in sensitivity to near half-saturating flashes under a higher dopamine concentration (250 μM).

**Figure 1.**
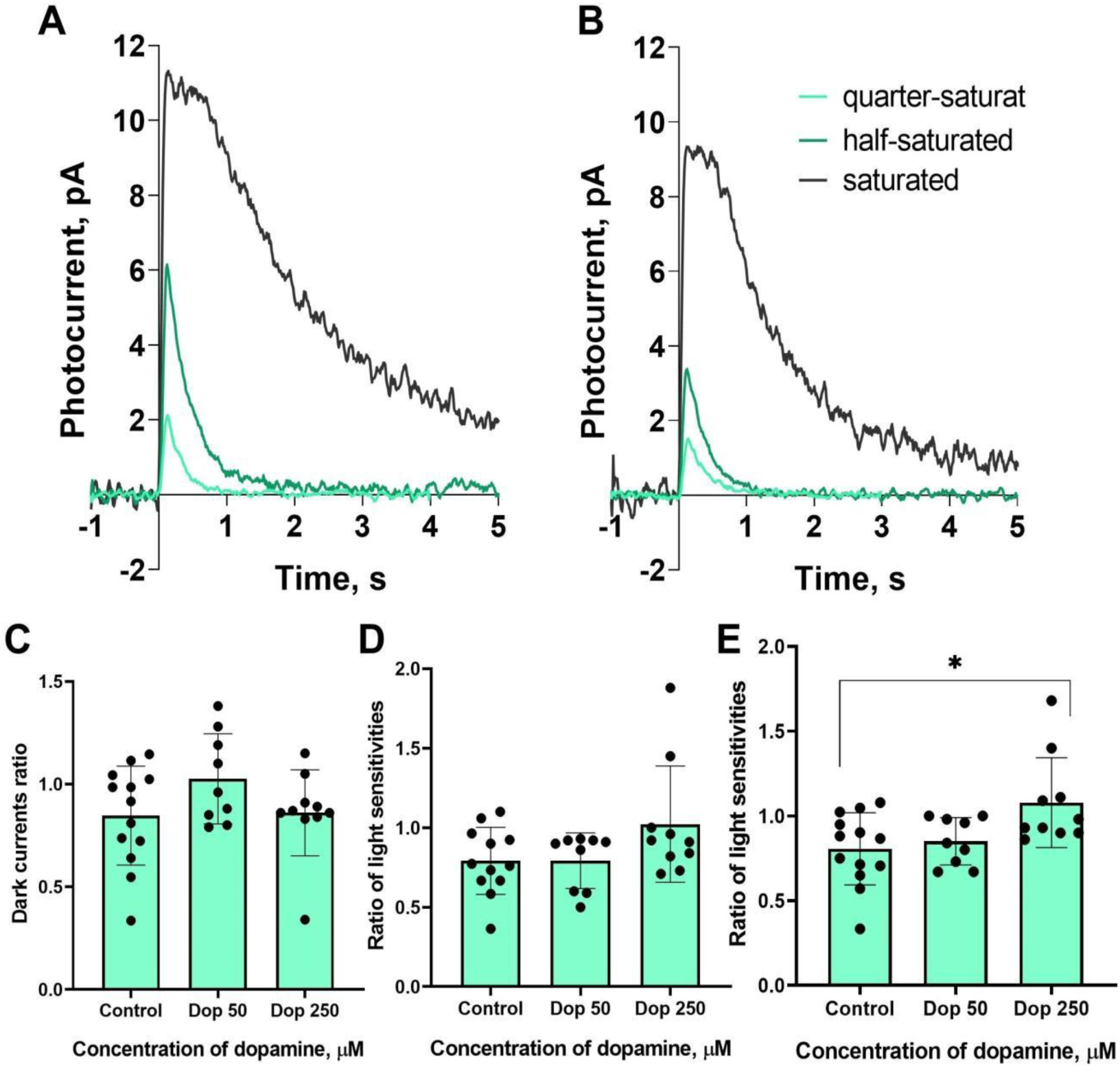
The effects of 50 and 250 μM dopamine on lamprey short photoreceptors. A typical set of responses of lamprey short photoreceptors to saturating, nearly half-saturating and quarter-saturating flashes in Ringer’s solution **(A)** and after 30 minutes of dopamine exposure **(B)**. Flash intensities are 1.03*10^4^ photons * μm^−2^ * s^−1^ (saturating), 246 photons * μm^−2^ * s^−1^ (near half-saturating) and 98 photons * μm^−2^ * s^−1^ (near quarter-saturating), λ_max_ = 521 nm. Figure 1(C) shows the effects of dopamine at both concentrations on the dark current, and **(D)** and (E) show the effects on light sensitivity to near quarter-saturating and near half-saturating flashes, respectively. * p < 0.05.

At first glance, there appeared to be no significant changes in the initial or falling phases of the photoresponses of the short photoreceptors during dopamine exposure. However, we decided to investigate this further. In order to analyse the behaviour of the initial phase of the response, we only needed to determine the relative changes in the steepness of the initial phase under two experimental conditions. While we did not provide a mathematical description of the biochemical mechanisms behind activation, we extracted the ratio of two activation rate parameters by adjusting the rising phases of these two photoresponses, as shown in Figure 2B.

**Figure 2.**
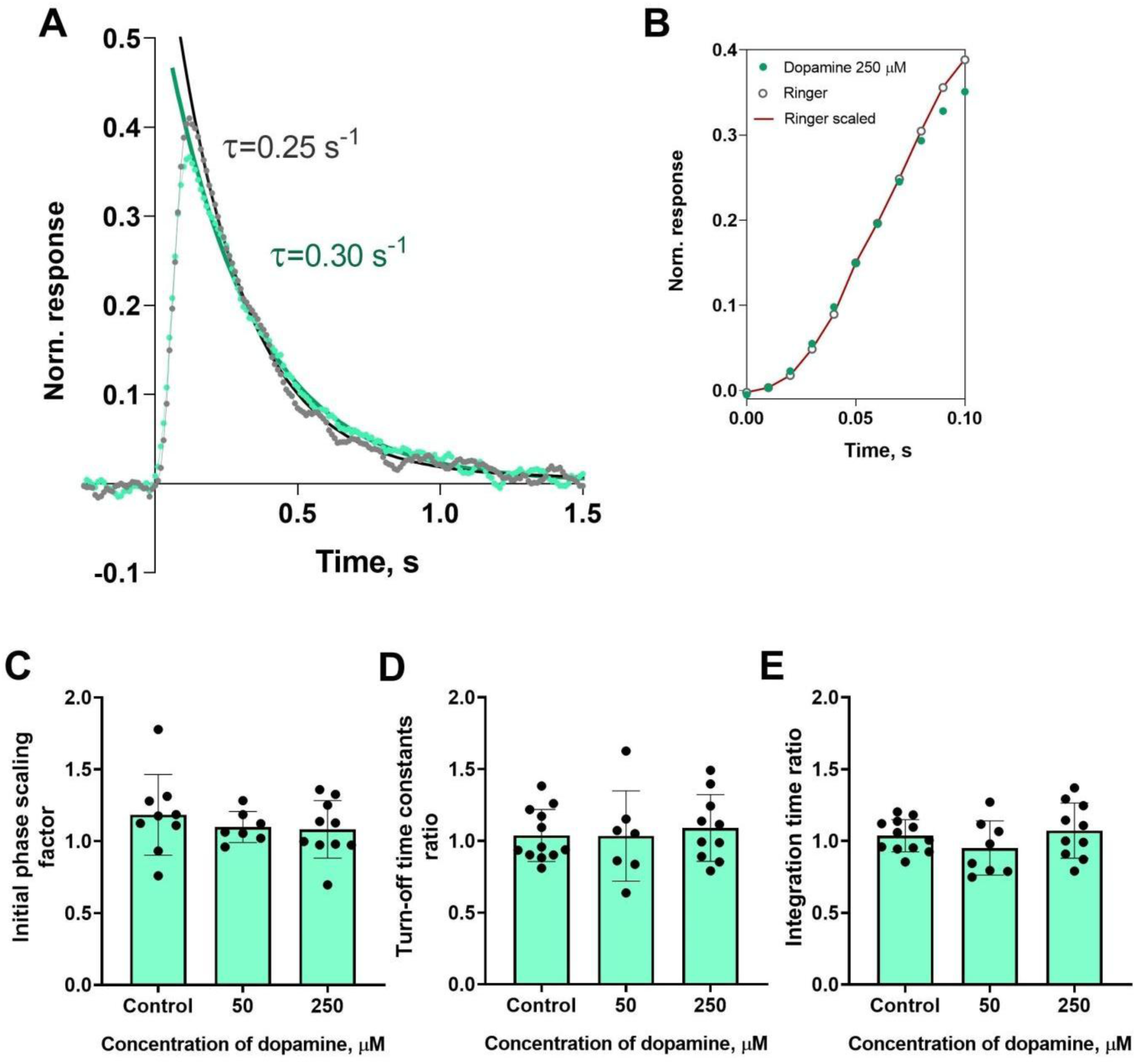
The effects of 50 and 250 μM dopamine on the photoresponse kinetics of lamprey short photoreceptors. **(A)** An example of normalized responses of lamprey long photoreceptors to near-half-saturating flashes in Ringer’s solution (grey curve and dots), and after 30 minutes of dopamine exposure (green curve and dots). The result of scaling the rising phase of these two photoresponses: the red curve shows the result of multiplying the initial phase of the response in Ringer’s solution by a scaling coefficient of 1.0. **(B)** Comparison of several response kinetic parameters: the scaling coefficient for the rising phase **(C)**, response recovery rates **(D)** and integration time **(E)** for responses recorded in Ringer’s solution and after 30 minutes of dopamine exposure.

To estimate the rate of the reactions underlying the falling phase of the photoresponse, we fitted the falling phase of near half-saturated responses with a single exponential function. We then extracted time constants τ, which were compared under two different conditions (see Figure 2A). We also compared the integration time of near half-saturated responses in Ringer’s solution and following the application of dopamine. Figure 2C, D and E show a summary of the results of the 30-minute dopamine exposure on the kinetic parameters and integration time of short photoreceptor responses in comparison with control experiments. Regardless of the tested concentration, there were no effects of dopamine on the kinetic parameters after 30 minutes of incubation. It should be noted that, at the intermediate time point of 20 minutes, we observed no effects of any dopamine concentrations on the dark current, light sensitivities, activation and falling phases, or the integration time of non-saturated responses (data shown in Fig.S1).

#### Long photoreceptors

The experimental protocol for testing the putative effects of dopamine on the long photoreceptors of lampreys was, in general, similar to those described above for the short photoreceptors. A set of responses, including saturated, near-half-saturated, and near-quarter-saturated ones, were recorded in Ringer’s solution, followed by recordings approximately 20 and 30 minutes after perfusion with Ringer’s solution containing 50 or 250 μM dopamine. We analyzed the same parameters for the long photoreceptors as we did for the short ones. Figure 3 shows a typical set of responses from lamprey long photoreceptors to saturating, near-half-saturating, and quarter-saturating flashes in Ringer’s solution and 30 minutes after dopamine exposure.

**Figure 3.**
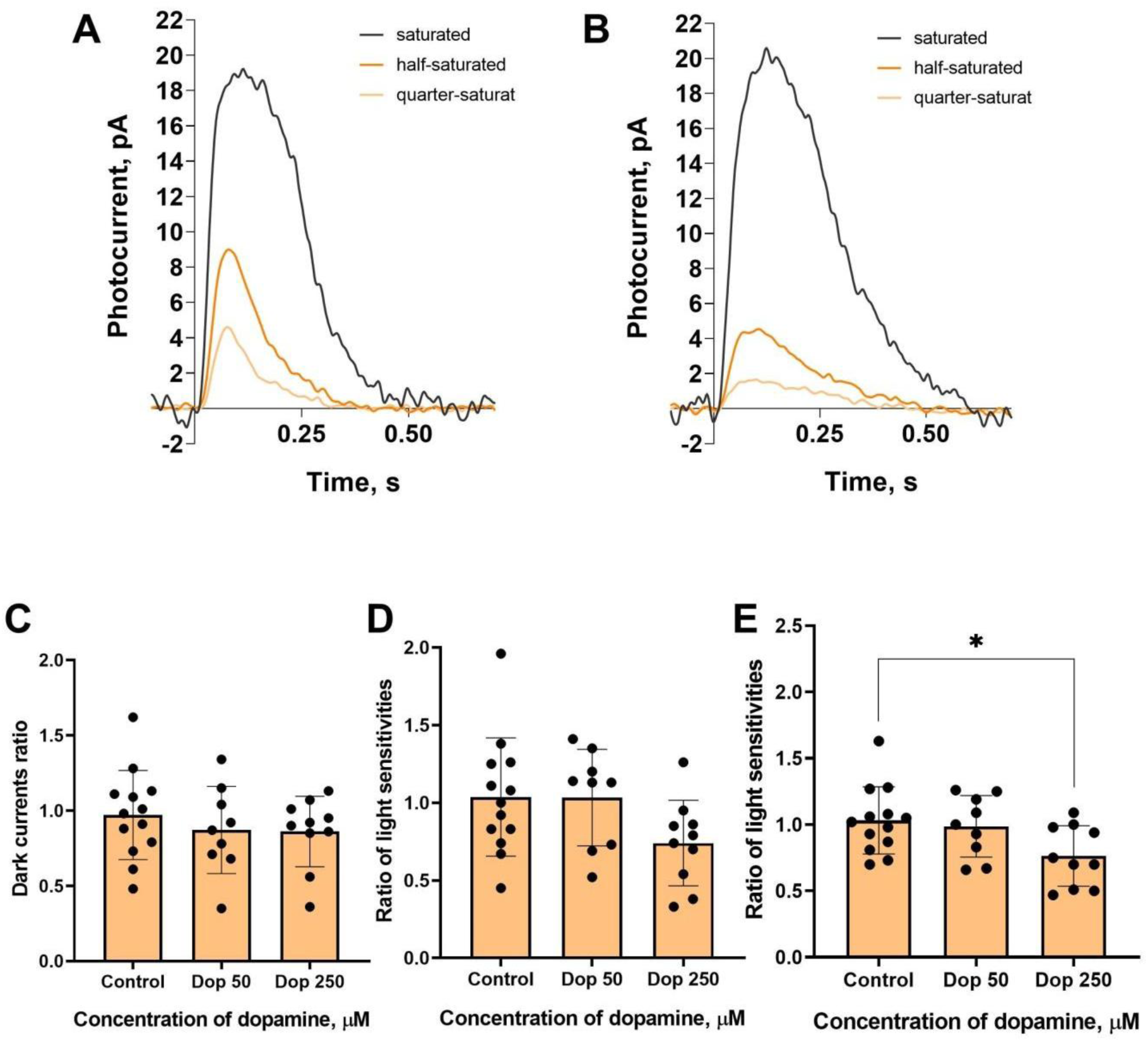
The effects of 50 and 250 μM dopamine on lamprey long photoreceptors. A typical set of responses from lamprey long photoreceptors to saturating, near-half-saturating and near-quarter-saturating flashes in Ringer’s solution **(A)**, and after approximately 30 minutes of dopamine exposure **(B)**. Flash intensities are 6.53*10^4^ photons * μm^−2^ * s^−1^ (saturating), 6.49*10^3^ photons * μm^−2^ * s^−1^ (near half-saturating) and 2.58*10^3^ photons * μm^−2^ * s^−1^ (near quarter-saturating), λ_max_ = 521 nm. Effects of dopamine at both concentrations on the dark current **(С)** and the light sensitivity to near quarter-saturating flashes **(D)** and half-saturating flashes **(E)**, respectively, for lamprey short photoreceptors. * p < 0.05.

At the 30-minute time point we observed a decrease in the light sensitivity under dopamine exposure at a concentration of 250 μM, which was statistically significant for the near half-saturated response. However, at the intermediate time point of a 20-minute incubation (see Fig. S2), the decrease in light sensitivity at 250 μM dopamine was not statistically significant.

Analysis of the response kinetics in the long photoreceptors revealed a statistically significant effect of dopamine at a high concentration on all tested parameters: dopamine slowed down both the activation and turn-off rates (Figures 4C and D), and as a result, increased the integration time of the non-saturated photoresponse (Figure 4E). These effects on the kinetic parameters were essentially the same at the intermediate time point of 20 minutes (see Figures S2D, E and F).

**Figure 4.**
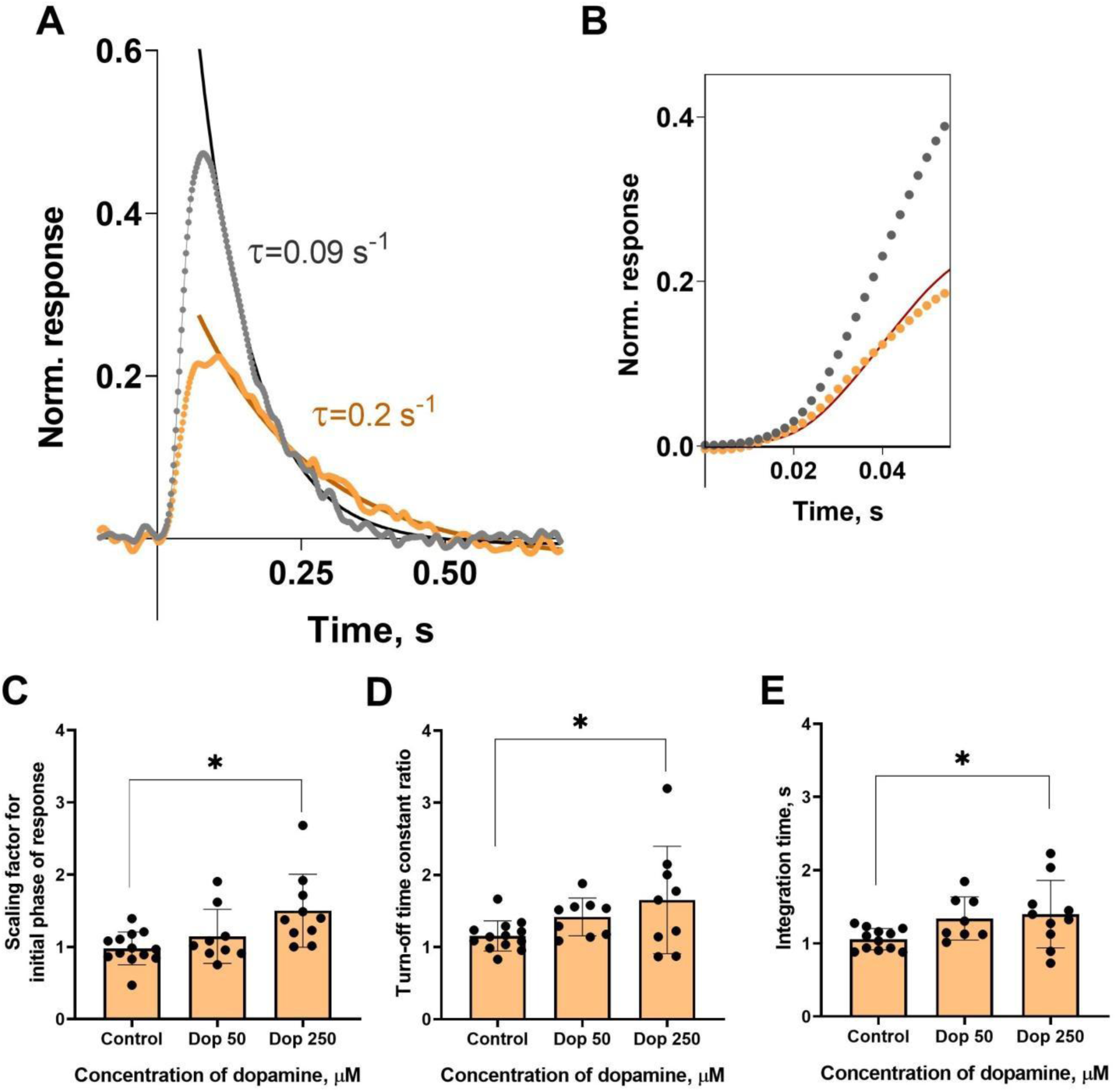
Effects of 50 and 250 μM dopamine on the photoresponse kinetics of lamprey long photoreceptors. **(A)** An example of normalized responses of lamprey long photoreceptors to near-half-saturating flashes in Ringer’s solution (grey curve and dots) and after 30 minutes of dopamine exposure (orange curve and dots). The result of scaling the rising phase of these two responses is shown by the red curve, which is the initial phase of the response in Ringer’s solution multiplied by a scaling coefficient of **0.54**. **(B)**. Comparison of several response kinetic parameters: the scaling coefficient for the rising phase **(C)**, the response recovery rates **(D)**, and the integration time **(E)** for responses recorded in Ringer’s solution and after 30 minutes of dopamine exposure. * p < 0.05.

### The lack of effect of increased cAMP levels on the photoresponse properties in both long and short photoreceptors of the river lamprey

We used the adenylcyclase activator forskolin to study the putative effects of increased intracellular cAMP levels on photoresponses of the river lamprey’s photoreceptors. Initially, photoresponses were recorded in Ringer’s solution. After that, the recordings were repeated after approximately 20- or 30-minute perfusion with forskolin-containing solution (10 μM). The same light stimulation protocol was used for both sets of recordings. The tested sets of responses included: 1) responses to a saturating flash; 2) responses to a half-saturating flash; and 3) responses to a quarter-saturating flash. Responses to non-saturating flashes were normalised to a dark current value at the corresponding time point, in order to compare response parameters in Ringer’s solution and after 20 or 30 minutes of forskolin application. The amplitudes of normalised non-saturating responses were used to measure photoreceptor sensitivity; the response kinetic parameters were also compared. The magnitude of the saturated responses under different conditions was compared as a measure of the dark current.

Forskolin did not affect the sensitivity or the kinetic parameters of photoresponses to non-saturating flashes in both short and long lamprey photoreceptors. Figures 5 and 6 show typical responses to a near-half-saturating flash before and after forskolin treatment. The responses before and after the exposure to forskolin look very similar. This was true for both the final (30 minutes) and intermediate (20 minutes) time points (see Fig. S3).

**Figure 5.**
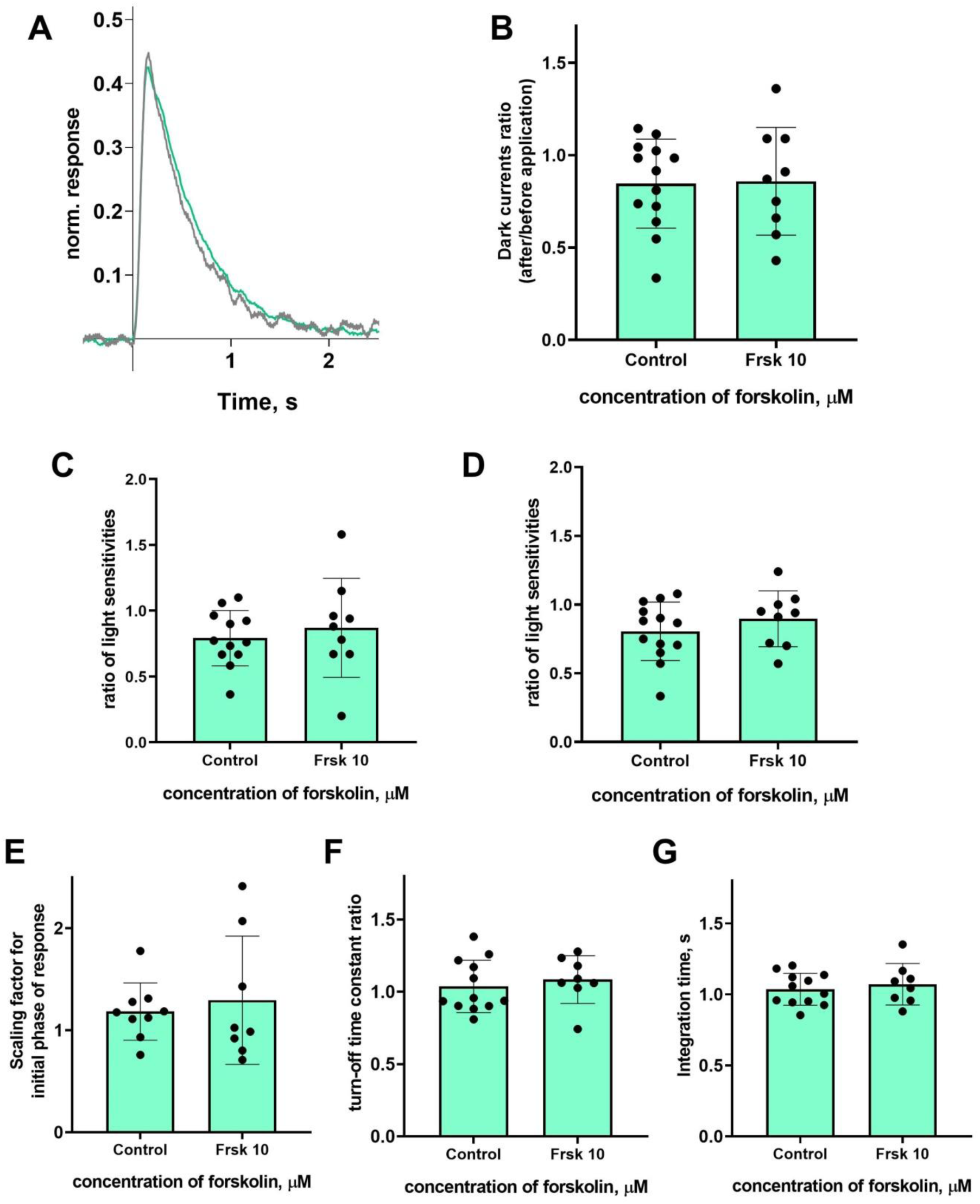
There is no effect of 10 μM forskolin on the photoresponses of short lamprey photoreceptors. **(A)** Responses of a representative short photoreceptor to brief, near-half-saturating flashes before (grey line) and after (green line) 30 minutes of forskolin application (each curve is an average of eight records obtained under the same conditions). Flash intensity: 61.8 photons * μm^−2^ * s^−1^, λ_max_ 521 nm. There is no difference in the dark current **(B)**, light sensitivity to near quarter-flashes **(C)**, or half-saturating flashes **(D)** for lamprey short photoreceptors after 30 minutes of forskolin exposure compared to the control experiments. There was no effect of forskolin on the kinetic parameters of the response: the scaling coefficient for the rising phase **(C)**, the response recovery rates **(D)** and the integration time **(E)** after 30 minutes of forskolin exposure.

**Figure 6.**
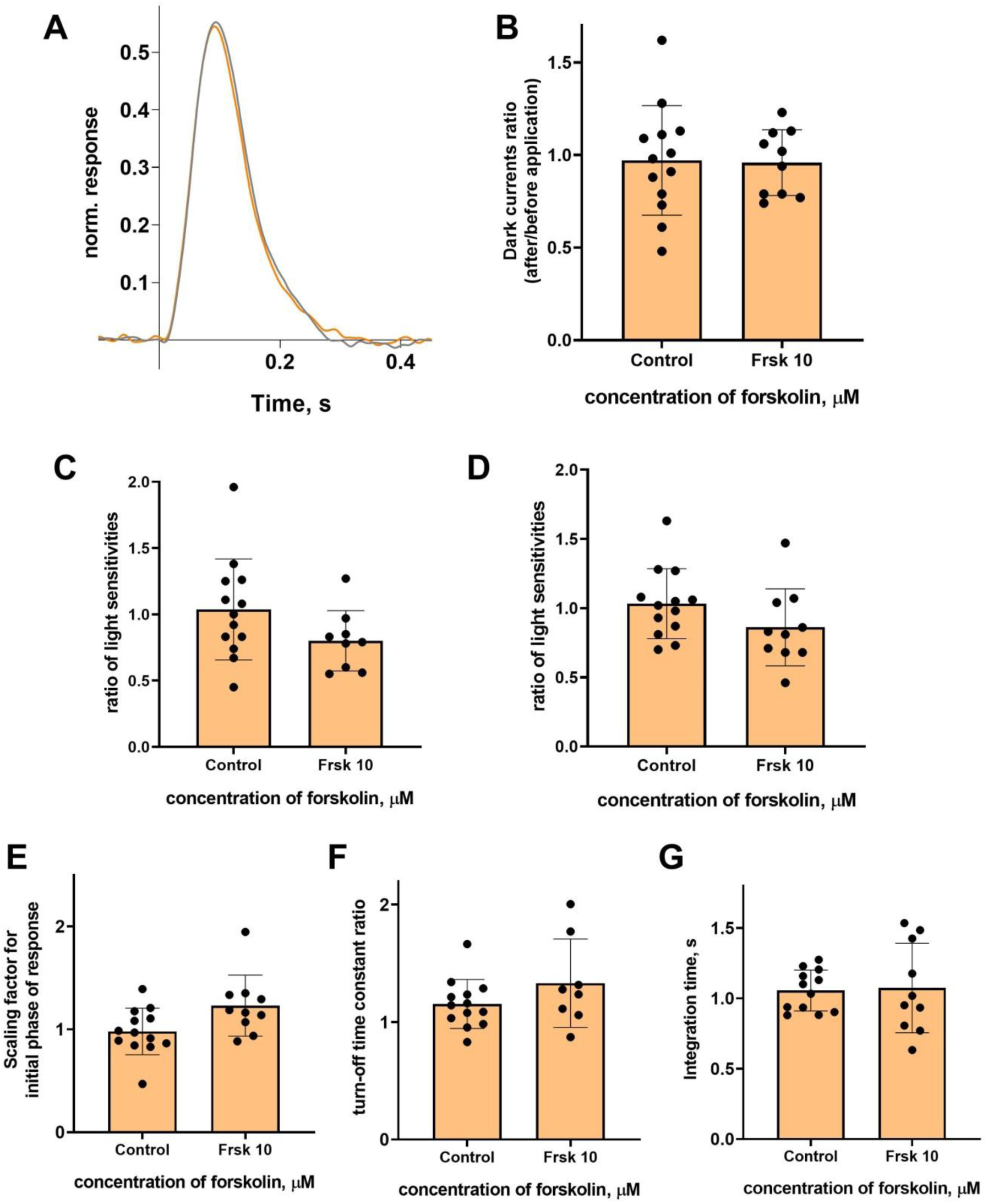
There is no effect of 10 μM forskolin on the photoresponses of long lamprey photoreceptors. **(A)** Responses of a representative long photoreceptor to brief, near-half-saturating flashes before and after 30 minutes of forskolin application (each curve is the average of 14 records obtained under the same conditions). Flash intensity: 409 photons * μm^−2^ * s^−1,^λ_max_ 521 nm. There is no difference in dark current **(B)**, light sensitivity to near quarter-saturating **(C)** and half-saturating **(D)** flashes for lamprey short photoreceptors after 30 minutes of forskolin exposure compared to control experiments. There was no effect of forskolin on the kinetic parameters of the response: the scaling coefficient for the rising phase **(C)**, the response recovery rates **(D)** and the integration time **(E)** after 30 minutes of forskolin exposure.

### The lack of dopamine effect on the photoresponse properties in fish green-sensitive cones

In previous subsections, we described the clear effects of dopamine application on the long photoreceptors of the river lamprey. We decided to compare the reactions of lamprey long photoreceptors to dopamine with the potential effects on typical vertebrate cones, since the two types of photoreceptors are functionally similar. To this end, we selected isolated green-sensitive cones from the fish *Carassius gibelio*. This type of photoreceptor was previously used in our long-term experiments and proved to be a stable and viable research subject.

In this experimental series, we only tested a higher concentration of dopamine (250 μM) on fish green-sensitive cones, which had an effect on lamprey long photoreceptors. The control experiments with the same experimental protocol were performed using only Ringer’s solution for green-sensitive fish cones in order to account for the putative changes in the main parameters of the photoresponses over time. We evaluated the same parameters as in the experiments involving lamprey long and short photoreceptors and took recordings at two time points: approximately 20 and 30 minutes after the start of the dopamine exposure.

Surprisingly, we observed no effect of dopamine on any of the measured parameters: dark current, light sensitivity, activation and turn-off kinetics, and integration time. A summary of the results of 30 minutes of dopamine exposure on fish cones is shown in Figure 7, and the results at the first time point (20 minutes after the start of dopamine exposure) are shown in the Fig. S5. There were no significant differences in any of the tested parameters at either time point compared to the control experiments.

**Figure 7.**
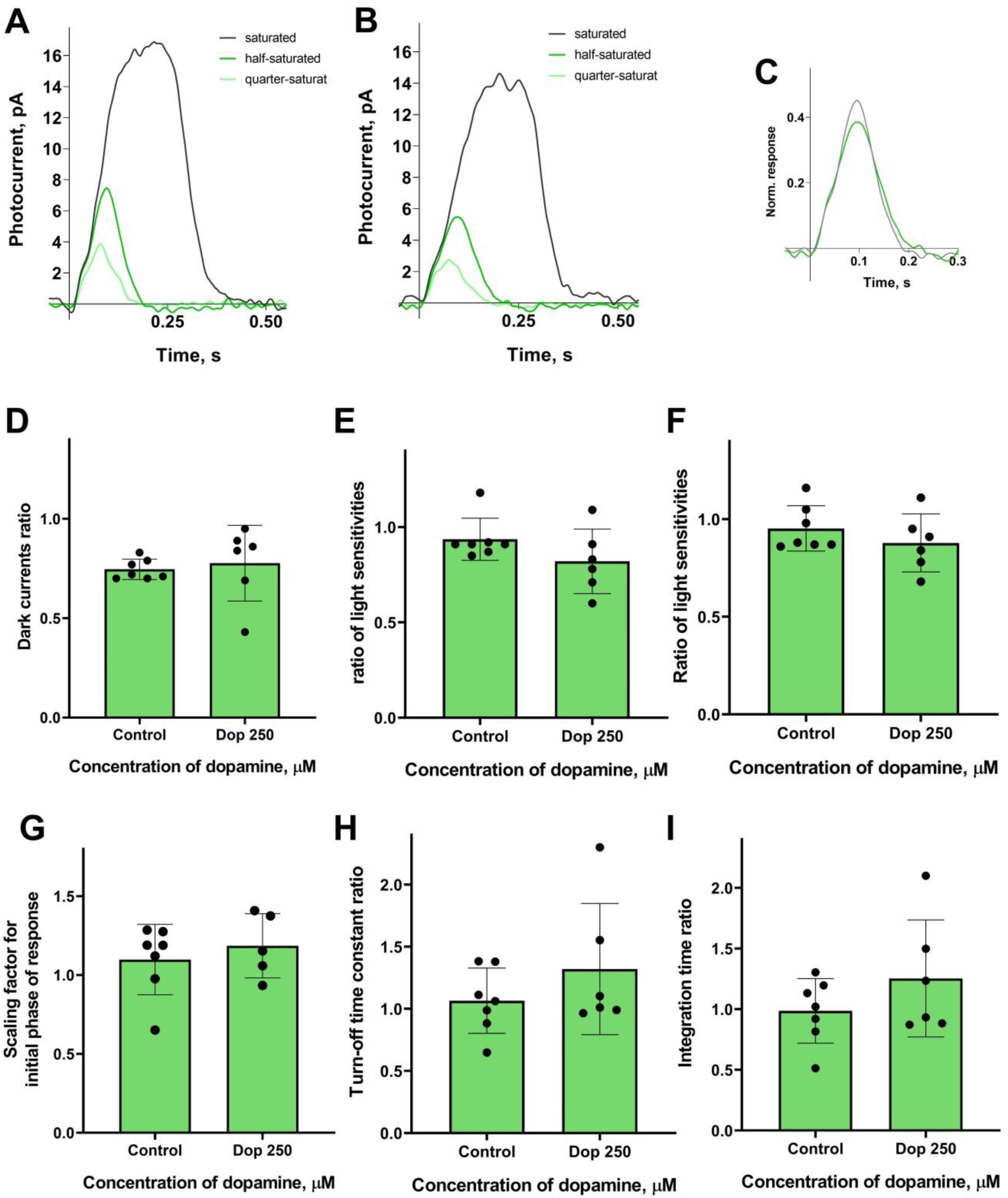
There is no effect of 250 μM dopamine on the green-sensitive cones of Carassius gibelio. A typical set of responses from a fish’s green-sensitive cone to saturating, near-quarter-saturating, and half-saturating flashes in Ringer’s solution **(A)** and after approximately 30 minutes of dopamine exposure **(B)**. **(C)** An example of normalized responses of fish cones to near-half-saturating flashes in Ringer’s solution (grey curve) and after 30 minutes of dopamine exposure (green curve). **(D)** and the light sensitivity to near half-saturating flashes **(E)** and quarter-saturating flashes **(F)** for fish cones. Comparison of several response kinetic parameters: **(G)** Scaling coefficient for the rising phase; **(H)** Response recovery rates; **(I)** Integration time for the responses recorded in Ringer’s solution and after 30 minutes of dopamine exposure.

### Immunoblotting and immunohistochemistry of D1RD and D2RD in the lamprey retina

Western blot results demonstrate the presence of D1RD-immunopositive bands in the lamprey retina corresponding to 55kDa and 100-130kDa, which is essentially similar to the distribution of D1RD immunopositive bands in the mouse retina (positive control, see Fig. S6A). This suggests the existence of monomeric and dimeric forms of the receptor, as well as its various glycosylated states (Jarvie et al., 1989; Luedtke et al., 1999).

Immunohistochemical analysis reveals a D1RD+ signal in various layers of the lamprey retina. A weak signal is detected in the photoreceptor layer (PL) in the outer segments of photoreceptors and a brighter signal is detected in the inner segments (Figure 8). A weak reaction is also detected in the layer corresponding to the OLM, in the ONL in the perikaryons of photoreceptor neurons as well as in the OPL. A more intense reaction is noted in the cell bodies and neuropil of the INL: in the cell bodies of large OHC cells (no reaction is noted in the bodies of small cells) and in the cell bodies of the IHC. An intense reaction is noted in the cell bodies of large INL cells localised in the OGCL (Figure 8). An intense reaction is also observed in the neuropil of the IPL layer and in the perikaryons and processes of ganglion cells located in the IGCL.

**Figure 8.**
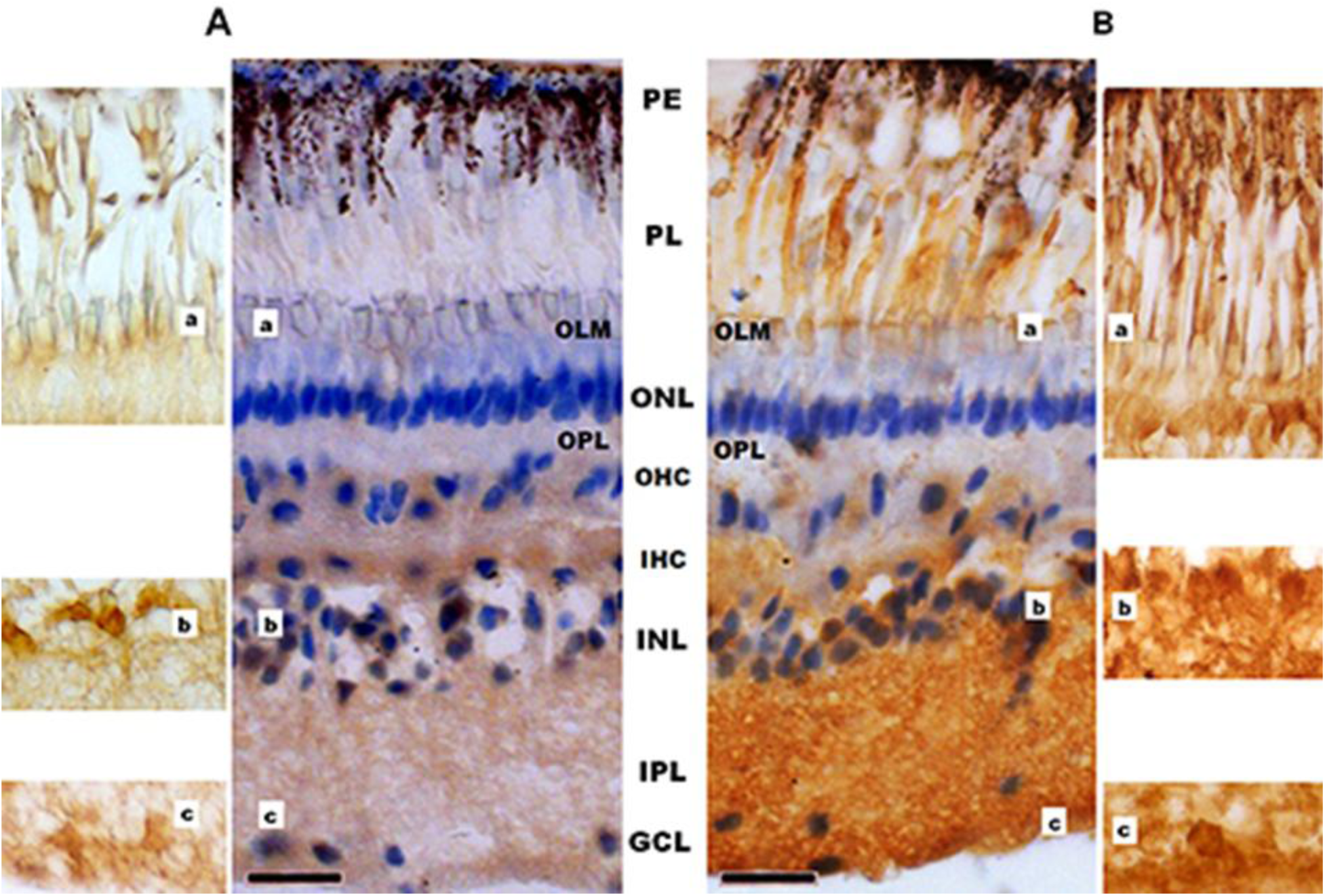
Cross-sections of the lamprey retina. Immunohistochemical reactions to D1RD (A) and D2RD (B). Counterstained with hematoxylin. Abbreviations (from Fritzsch and Collin, 1990): PE: pigment epithelium; PL: photoreceptor layer; OLM: outer limiting membrane; ONL: outer nuclear layer; OPL: outer plexiform layer; OHC: outer horizontal cells; IHC: inner horizontal cells; INL: inner nuclear layer; IPL: inner plexiform layer; GCL: ganglion cell layer. Scale bars: 25 µm. Zoomed-in images from the corresponding areas without haematoxylin staining.

Western blot results demonstrate the presence of immunopositive bands for D2R in the lamprey retina that correspond to 100kDa (as in the mouse retina), as well as additional bands above 55kDa that are not detected in the mouse retina. This may indicate greater dimerization of D2 receptors in the mouse retina than in the lamprey retina, as well as the separate detection of S and L isoforms of D2RD in the lamprey retina (see Fig. S6-B).

Immunohistochemical analysis reveals an intense reaction to D2RD in various retinal layers (Fig. 8). This reaction is present in the outer segments of photoreceptors (PL), at the OLM, and a weak reaction is noted in the perikaryons of photoreceptor neurons (ONL) and OPL. A D2RD+ reaction is also observed in the perikaryons and neuropil of the INL and IPL layers. The most intense reaction is seen in the perikaryons of large cells (OGCL and IGCL) (see Fig. 8).

### Immunoblotting and immunohistochemistry of D1RD and D2RD in the retina of Carassius gibelio

Western blot results show a distribution of D1RD immunopositive bands in the fish retina (50kDa, 100kDa and 130kDa regions) and additional bands (Fig. S8-A), similar to the distribution in the mouse retina.

Immunohistochemical analysis of D1RD distribution in the Carassius retina shows no reaction in the PL. In the ONL, the reaction is only detected in the perikaryons of cells located in the lower part of this layer (Figure 9). A weak reaction is detected in the OPL. In the INL, the reaction is detected in the perikaryons of cells, but the intensity varies between cells in this layer. A positive reaction to D1RD is evident in the IPL, particularly in the cells and processes of the GCL. The prevalence of D1RD-positive cells in the plexiform layers is consistent with previous immunohistochemical studies of fish retinas (Behrens and Wagner, 1995; Mora-Ferrer et al., 1999).

**Figure 9.**
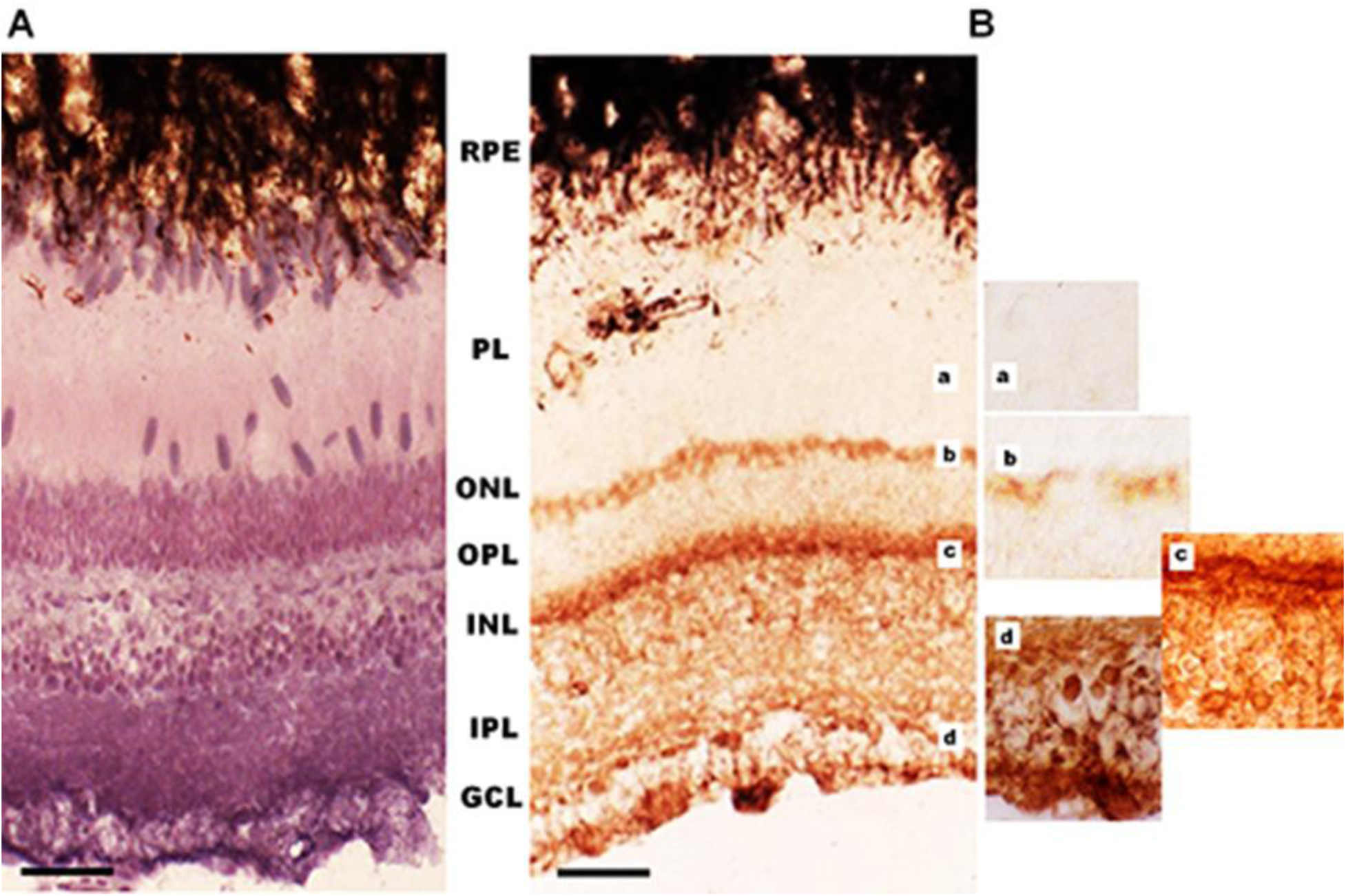
Cross-sections of the retina of *Carassius gibelio*: haematoxylin staining (A) and immunohistochemistry for D1RD (B). Abbreviations (from Beraudi et al., 2007): RPE: pigment epithelium; PL: photoreceptor layer; ONL: outer nuclear layer; OPL: outer plexiform layer; INL: inner nuclear layer; IPL: inner plexiform layer; GCL: ganglion cell layer. Scale bars: 25 µm. Zoomed-in images of the corresponding areas.

Western blot analysis of the retina of *Carassius gibelio* revealed extremely weak D2RD immunopositive bands at 100kDa (Fig. S8B) compared to the mouse retina, as well as more intense bands in the lower mass range (lower part of the gel).

Immunohistochemical analysis of D2RD distribution in the Carassius retina (Fig. 10) shows a weak D2RD+ reaction in the PL. In the ONL, the reaction is detected in the perikaryons of photoreceptors located in the lower part of this layer. A weak D2RD+ reaction is detected in the OPL, whereas a more intense and uniform reaction is noted in the perikaryons of INL cells. An intense D2RD+ reaction is also noted in the perikaryons and processes of GCL cells. A higher abundance of D2-like receptors than D1-like receptors has been observed previously (Wagner et al., 1993), although the authors could not observe specific staining in the retina of Carassius species with the available antiserums.

**Figure 10.**
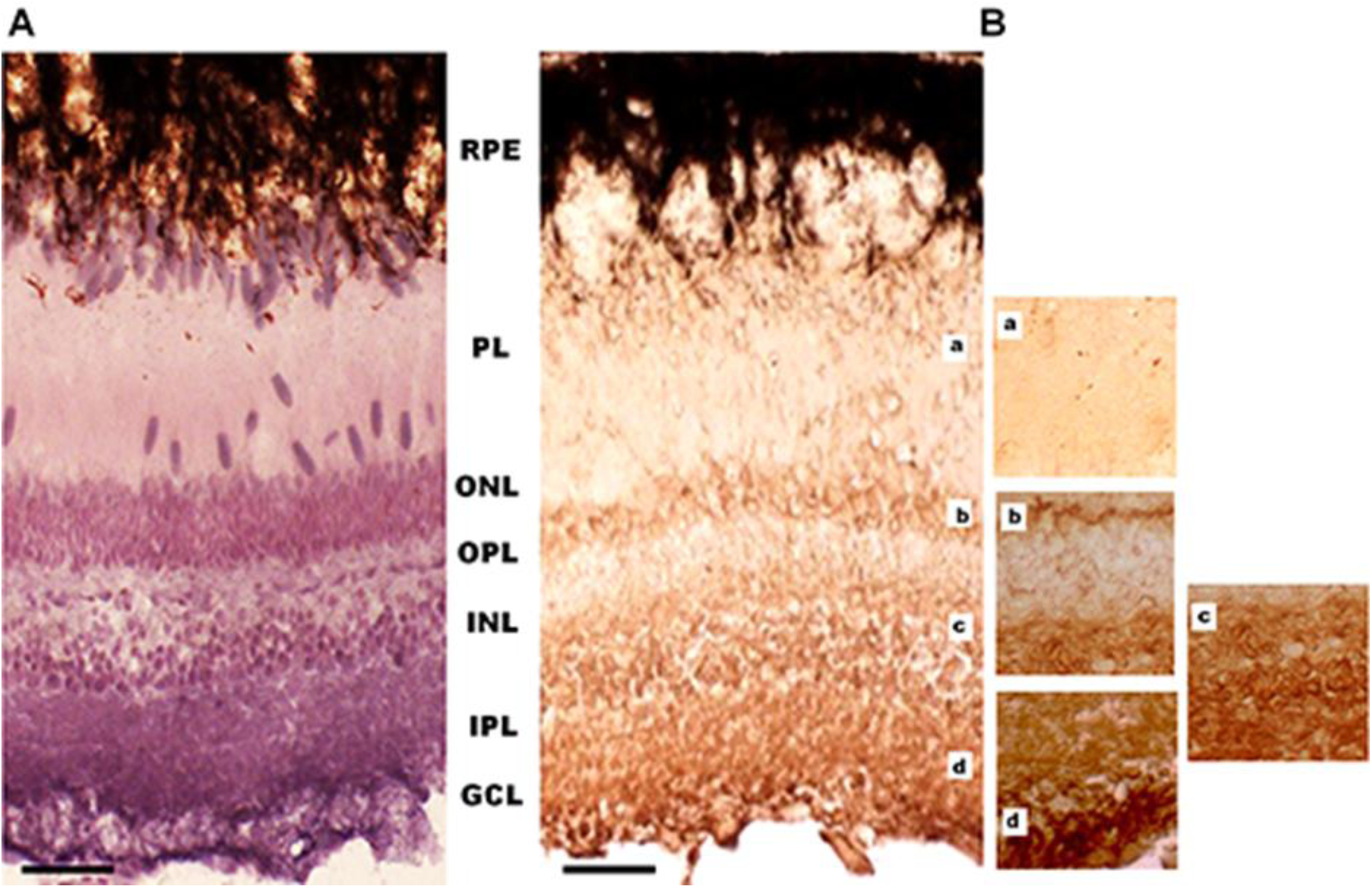
Cross-sections of the retina of Carassius gibelio: haematoxylin staining (A) and immunohistochemistry for D2AR (B). Abbreviations (from Beraudi et al., 2007): RPE: pigment epithelium; PL: photoreceptor layer; ONL: outer nuclear layer; OPL: outer plexiform layer; INL: inner nuclear layer; IPL: inner plexiform layer; GCL: ganglion cell layer. Scale bars: 25 µm. Zoomed-in images of the corresponding areas.

### Immunoblotting and immunohistochemistry of D1RD and D2RD in the retina of marsh frog

The Western blot results for the frog retina show a distribution of immunopositive dopamine D1 receptor bands in the 50, 100 and 130kDa regions, as well as the presence of additional bands similar to those in the mouse retina (Fig. S8-A). These data clearly indicate the presence of both monomeric and dimeric forms of the receptors, as well as their modifications.

The immunohistochemical results for the frog retina (Fig. 11) show a D1RD+ reaction in the outer segments of photoreceptors in the PL (mainly in the lower part, above the nuclei), as well as in the perikaryons of photoreceptors in the ONL. A D1RD+ reaction is clearly detected in the OPL and in some perikaryons located in the INL. An intense D1RD+ reaction is evident in the IPL, in the perikaryons, and in the processes of GCL cells. The detection of D1-like receptors in the retinal plexiform layers of frogs is consistent with earlier studies, while their potential localisation to photoreceptors remains a matter of debate (Behrens and Wagner, 1995; Zhekova and Vitanova, 2016).

**Figure 11.**
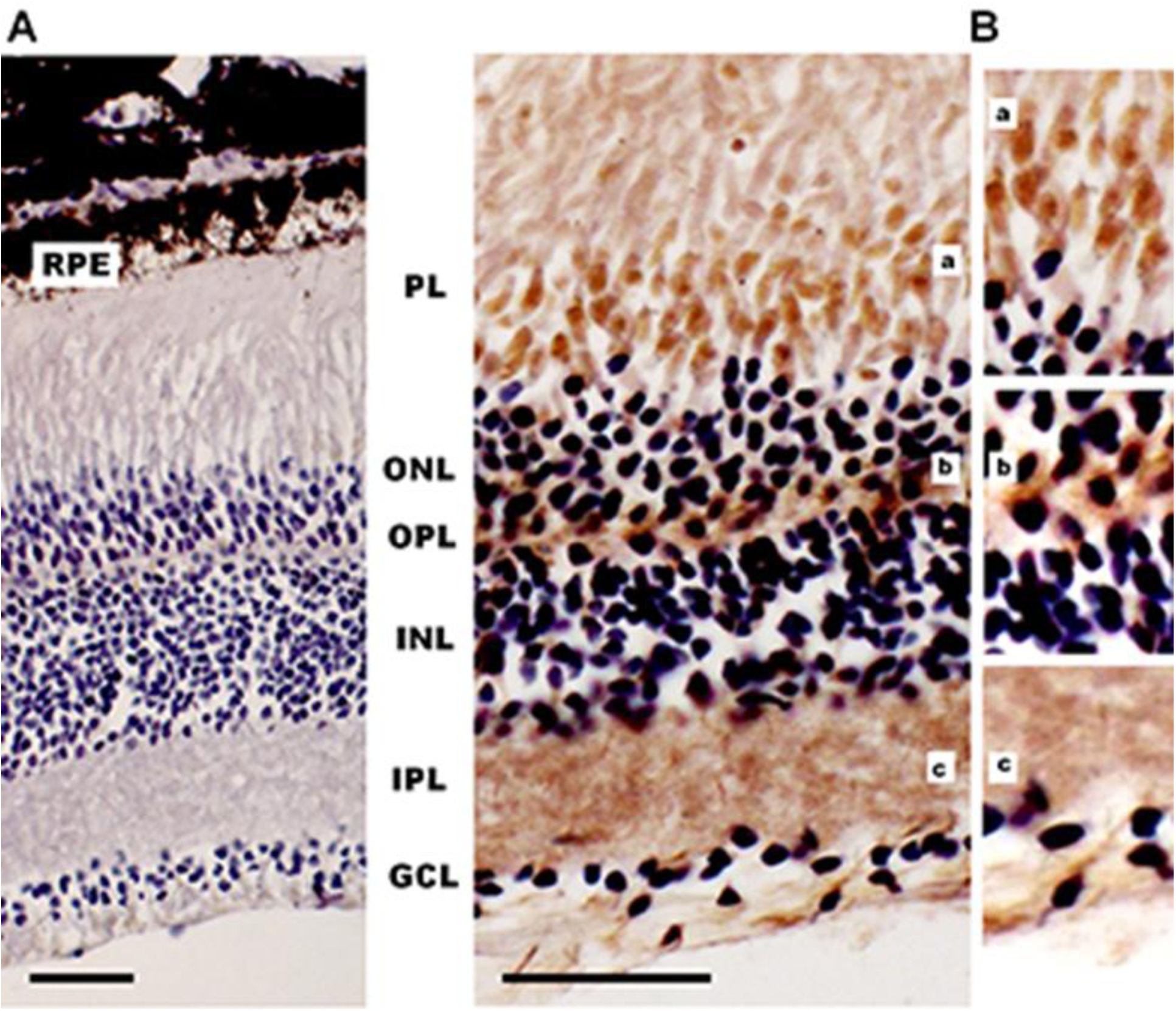
Cross-sections of the retina of a marsh frog *(Pelophylax ridibundus)*: hematoxylin staining (A) and immunohistochemistry for D1RD (B). Abbreviations (from Beraudi et al., 2007): RPE: pigment epithelium; PL: photoreceptor layer; ONL: outer nuclear layer; OPL: outer plexiform layer; INL: inner nuclear layer; IPL: inner plexiform layer; GCL: ganglion cell layer. Scale bars: 25 µm. Zoomed-in images of the corresponding areas.

Western blot analysis of the frog retina reveals D2RD immunopositive bands around 100kDa, which are similar to those in the mouse retina. This demonstrates that dimeric forms of D2RD are formed in both species (see Fig. S8-B).

Immunohistochemical analysis of the frog retina (Figure 12) reveals a D2RD+ reaction in the lower sublayer of the PL, but this reaction is only detected in the central part of the retina. In the peripheral retina, the PL does not react to D2RD. The D2RD+ reaction is clearly detected in the perikaryons of photoreceptors (ONL) and in the neuropil of the OPL, as well as in the perikaryons of individual neurons of the INL. An intense D2RD+ reaction is also detected in the IPL, as well as in the perikaryons and processes of ganglion neurons (GCL). This D2-like expression pattern matches earlier reports on the Xenopus retina (Wagner et al., 2003).

**Figure 12.**
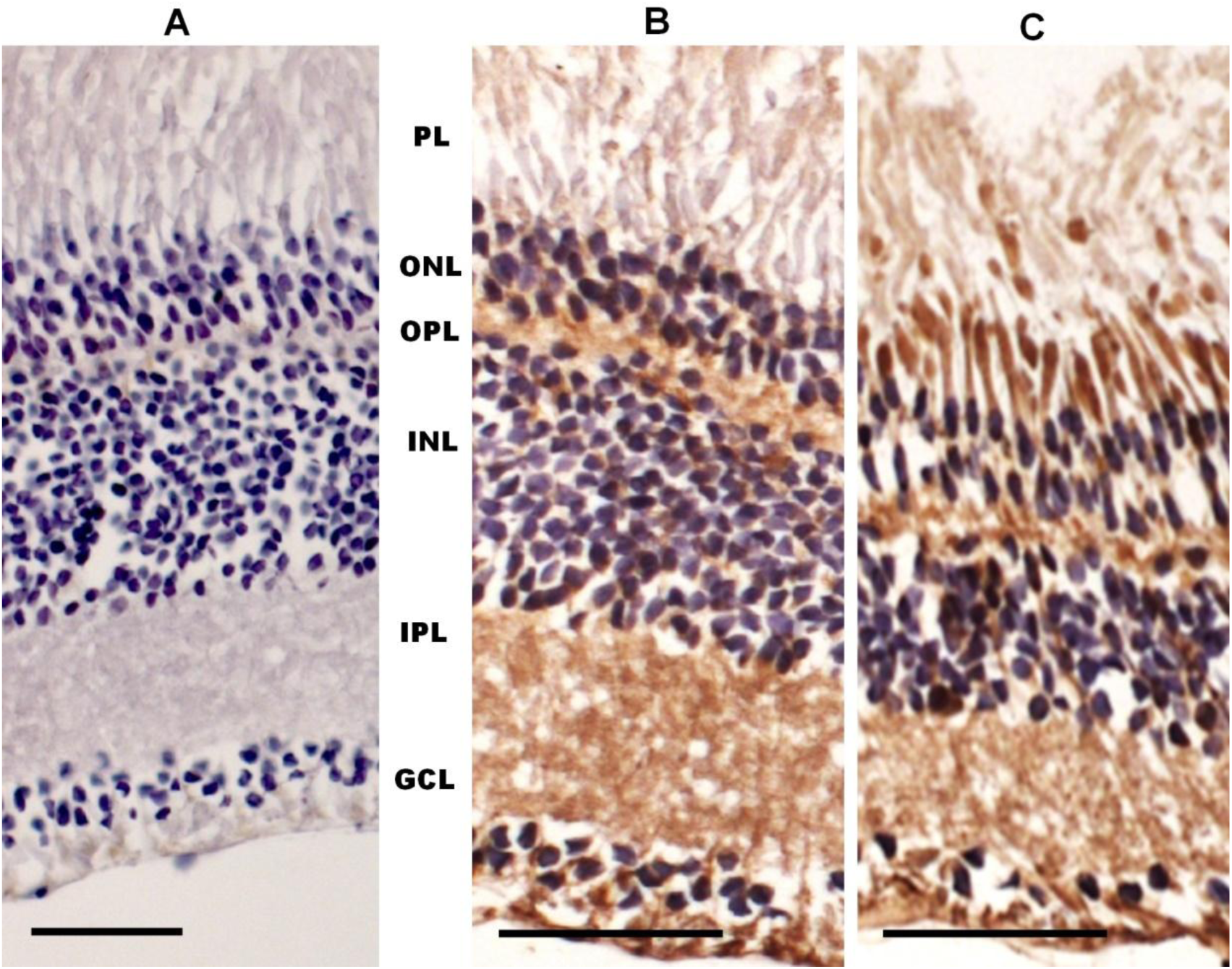
Cross-sections of the retina of a marsh frog: haematoxylin staining (A), and immunohistochemistry for the peripheral (B) and central (C) parts of the retina. Abbreviations (from Beraudi et al., 2007): RPE: pigment epithelium; PL: photoreceptor layer; ONL: outer nuclear layer; OPL: outer plexiform layer; INL: inner nuclear layer; IPL: inner plexiform layer; GCL: ganglion cell layer. Scale bars: 25 µm. Zoomed-in images of the corresponding areas.

## Discussion

Our results unequivocally demonstrate the existence of a regulatory mechanism involving dopamine that modulates the sensitivity and kinetics of lamprey photoreceptors in response to light. However, the nature of this regulation differs markedly from that observed for the rods and cones of jawed vertebrates. Notably, the involvement of intracellular cAMP concentration in regulating the phototransduction cascade in lampreys is either absent or only can be observed when its concentration would be artificially decreased.

### Lamprey circadian rhythms and dopamine

Northern hemisphere lampreys exhibit relatively primitive light-dependent behavior, primarily manifested by negative phototaxis. In the larval stage, ammocoetes, and during seasonal upstream migration in rivers, lampreys only actively move at night and utilise habitats during the day (Hardisty, 2013; Zvezdin et al., 2019). During their active feeding period at sea, adult lampreys synchronize their locomotor activity with the daily vertical movements of prey fish in the pelagic zone (Maitland, 2003). The circadian rhythm of lamprey locomotor activity is regulated by an oscillator in the well-developed pineal eye (Cole and Youson, 1982; Tamotsu and Morita, 1986) and persists even after the lateral eyes have been surgically removed (Morita et al., 1992). Nevertheless, the isolated lamprey retina has been shown to possess its own circadian oscillator that controls the expression of melatonin by photoreceptors (Menaker and Tosini, 1996). However, the involvement of retinal dopamine in the regulation of lamprey locomotor activity is doubtful, since dopaminergic amacrine cells only appear at a relatively late stage in the lamprey’s ontogeny. In the retinas of ammocoetes, which exhibit a clear diurnal rhythm of locomotor activity, these cells only appear from stages close to metamorphosis into adult lampreys (Villar-Cerviño et al., 2006; Abalo et al., 2008).

The role of dopamine in the lamprey retina may not be directly linked to visual perception, as it appears to be less dependent on the brain than in higher vertebrates. For instance, the light pupillary reflex in lampreys is not mediated by the brain (Jiménez-López et al., 2024) and is instead fully controlled by a neural circuit within the eye. This circuit is most likely triggered by light-sensitive retinal cells containing melanopsin (Morshedian et al., 2021). Rather, a supporting role might be played by dopamine by adjusting the sensitivity of the visual system to different light levels. This hypothesis is supported by the expression of dopamine receptors in many retinal cell types. In teleosts and amphibians, the pineal organ also represents an independent circadian oscillator (Harada et al., 1998; Saha et al., 2019). However, in the retina, dopamine levels change cyclically (increasing during the day and decreasing at night) in an independent manner, primarily in response to changes in illumination and, to a lesser extent, to the activity of the internal retinal oscillator (Ribelayga and Mangel, 2003). The regulation of various aspects of photoreceptor physiology by dopamine has been convincingly demonstrated (see Popova, 2020 for a review). The role of dopamine in directly regulating the phototransduction cascade has only recently been evaluated (Nikolaeva et al., 2019). As dopamine significantly impacts direct signal transmission between photoreceptors via gap junctions, its undistorted impact on photoresponse sensitivity and kinetics can only be investigated through recordings from isolated cells or small retinal fragments.

### Dopamine signalling pathways affecting the phototransduction cascade

Vertebrate photoreceptors are traditionally considered to express exclusively D2-like dopamine receptors (Mureșan and Besharse, 1993; Vuvani et al., 1993; Wagner et al., 1993), though some authors have suggested the presence of D1-like receptors (Bjelke et al., 1996; Mora-Ferrer et al., 1999). D1- and D2-like dopamine receptors are two classes that demonstrate opposite effects on intracellular cAMP levels. D1-like receptors (subtypes D1 and D5) increase the activity of adenylyl cyclase (AC) via the Gs pathway, whereas D2-like receptors (subtypes D2, D3 and D4) decrease AC activity via the inhibitory Gi pathway (Nguyen-Legros et al., 1999; Vallone et al., 2000). Despite having fewer subtypes, lampreys have been shown to possess receptors from both the D1- and D2-like families (Robertson et al., 2012; Yamamoto et al., 2013). Another signalling pathway triggered by dopamine receptors that is unrelated to ACs is based on the activation of Gq, which regulates phospholipase C (PLC) activity and leads to the massive release of calcium ions from intracellular depots (Missale et al., 1998; Sahu et al., 2009). Thus, cAMP is one of the key mediators in the regulation of intracellular processes by dopamine. Its main target, protein kinase A (PKA), is activated when the level of cAMP increases and is inhibited when it decreases (Beaulieu and Gainetdinov, 2011). Several key proteins in the phototransduction cascade are PKA substrates: guanylate cyclase (Wolbring and Schnetkamp, 1996) and GC-activating proteins (Peshenko et al., 2004); opsin kinases GRK7 and GRK1 (Osawa et al., 2008, 2011) and phosducin (Pagh-Roehl et al., 1995; Willardson et al., 1996). Changes in retinal cAMP levels follow a circadian rhythm, controlling the rate of melatonin synthesis in photoreceptors (Iuvone and Besharse, 1986; Pierce et al., 1989; Cohen and Blazynski, 1990). This, in turn, affects the release of dopamine by amacrine cells in the inner retina. Consequently, a feedback loop via dopamine receptors controls the day/night cycle of retinal function (Nir et al., 2002; Tosini et al., 2008).

Frog rod photoresponses are unaffected by dopamine concentrations of up to 50 μM when applied to the inner segment. However, application to the outer segment produced significant modulation: sensitivity decreased mainly due to a slower activation rate of the cascade, and to a lesser extent due to faster inactivation (Nikolaeva et al., 2019). We have demonstrated that lamprey short photoreceptor responses remain unaffected by the application of 50 μM dopamine to the inner segment. However, a slight increase in sensitivity was observed when the concentration was increased to 250 μM.

Our immunohistochemical analysis in the present study showed that frog rods express both D1 and D2 receptor subunits, with a higher abundance in the inner segment than in the outer segment. Our previous electrophysiological study using selective agonists of different types of dopamine receptors indicated that the observed effect of dopamine on frog rods is most likely mediated by heteromeric D1–D2 receptors in the outer segment and not by homomeric D1-like or D2-like receptors (Nikolaeva et al., 2019). Furthermore, decreased sensitivity cannot be fully explained by cAMP-mediated signalling pathways and must also involve the regulation of intracellular calcium. Application of forskolin (2 μM) significantly increases rod sensitivity by slowing down shut-down processes, but does not activate phototransduction (Astakhova et al., 2012; 2017). It is potentially the case that D1 and D2 receptors in the inner segment do not form heteromeric complexes, or it may be that calcium level modulation only has an effect when it occurs in the outer segment.

Our immunohistochemical results indicate that lamprey short photoreceptors predominantly express D2-like subunits in the outer segment and show a weak D1-like signal in the inner segment. The slight increase in sensitivity upon application of 250 μM dopamine could be explained by its action via D1-like receptors. However, experiments involving forskolin demonstrate the complete insensitivity of lamprey short photoreceptors to elevated cAMP levels. Given the dense distribution of D2-like receptors in the outer segment, it is likely that dopamine can modulate the photoresponses of short photoreceptors at lower concentrations. This modulation could be achieved through changes in calcium levels, a decrease in intracellular cAMP or direct interaction between dopamine receptors and certain ion channels. (Kiselevsky et al., 2008a; 2008b).

Similarly to frog rods, fish cones were not sensitive to the application of dopamine to the inner segment, even at a concentration of 250 μM. By contrast, the responses of lamprey long photoreceptors were significantly modulated. The on- and off-cascade processes slowed simultaneously, resulting in an increase in integration time; however, sensitivity to brief flashes of light remained unchanged. Interestingly, a similar effect was observed in the red- and green-sensitive cones of fish following the application of forskolin (Sitnikova et al., 2021), which suggests that, in lamprey long photoreceptors, dopamine also mediates an increase in intracellular cAMP concentration. However, both short and long lamprey photoreceptors were insensitive to forskolin, even at higher concentrations (10 μM vs. 2 μM), rendering this assumption untenable.

The immunohistochemical analysis of fish photoreceptors presented here shows that there is weak expression of the D2-like receptor and no expression of the D1-like receptor in the outer and inner segments. This is consistent with earlier reports (Wagner et al., 1993; Behrens, Wagner, 1995; Mora-Ferrer et al., 1999). Lamprey long photoreceptors (photoreceptors most similar to cones in higher vertebrates) exhibit a dopamine receptor expression profile similar to that of short photoreceptors, with weak D1 receptor expression in the inner segment and a dense distribution of D2 receptors in the outer segment. The relatively low sensitivity of lamprey photoreceptors to dopamine is also consistent with dopamine acting via D1-like rather than D2-like receptors (Kebabian & Calne, 1979; Missale et al., 1998). While dopamine can increase intracellular cAMP levels via D1-like receptor downstream signalling, the insensitivity of long photoreceptors to forskolin suggests the presence of alternative signalling pathways that affect the phototransduction cascade. In particular, the mechanism of slowing down shut-down associated with the modulation of cone-specific opsin kinase GRK7 activity via PKA is ruled out for lamprey receptors (Chrispell et al., 2022).

### Physiological role of dopamine regulation: lamprey vs gnathostomes

The regulation of the gnathostomes’ phototransduction cascade by dopamine is far from fully understood in terms of its signalling pathways and the exact participants involved. However, the available data (summarized in Table 1) allow us to draw several general conclusions: (1) Dopamine has no effect when acting on receptors of the inner segment; (2) When acting on the outer segment of rods, dopamine decreases their sensitivity during the daytime; (3) An increase in intracellular cAMP concentration, which corresponds to night-time, increases the sensitivity of rods and slows the kinetics of cone photoresponses. It seems that, through dopamine-mediated regulation, rods gain an additional mechanism to adjust their sensitivity to high or low light conditions. The decrease in the activity of cone adaptation mechanisms at night excludes their participation in the visual process but may put them into metabolic economy mode (Sitnikova et al., 2021).

**Table 1.**
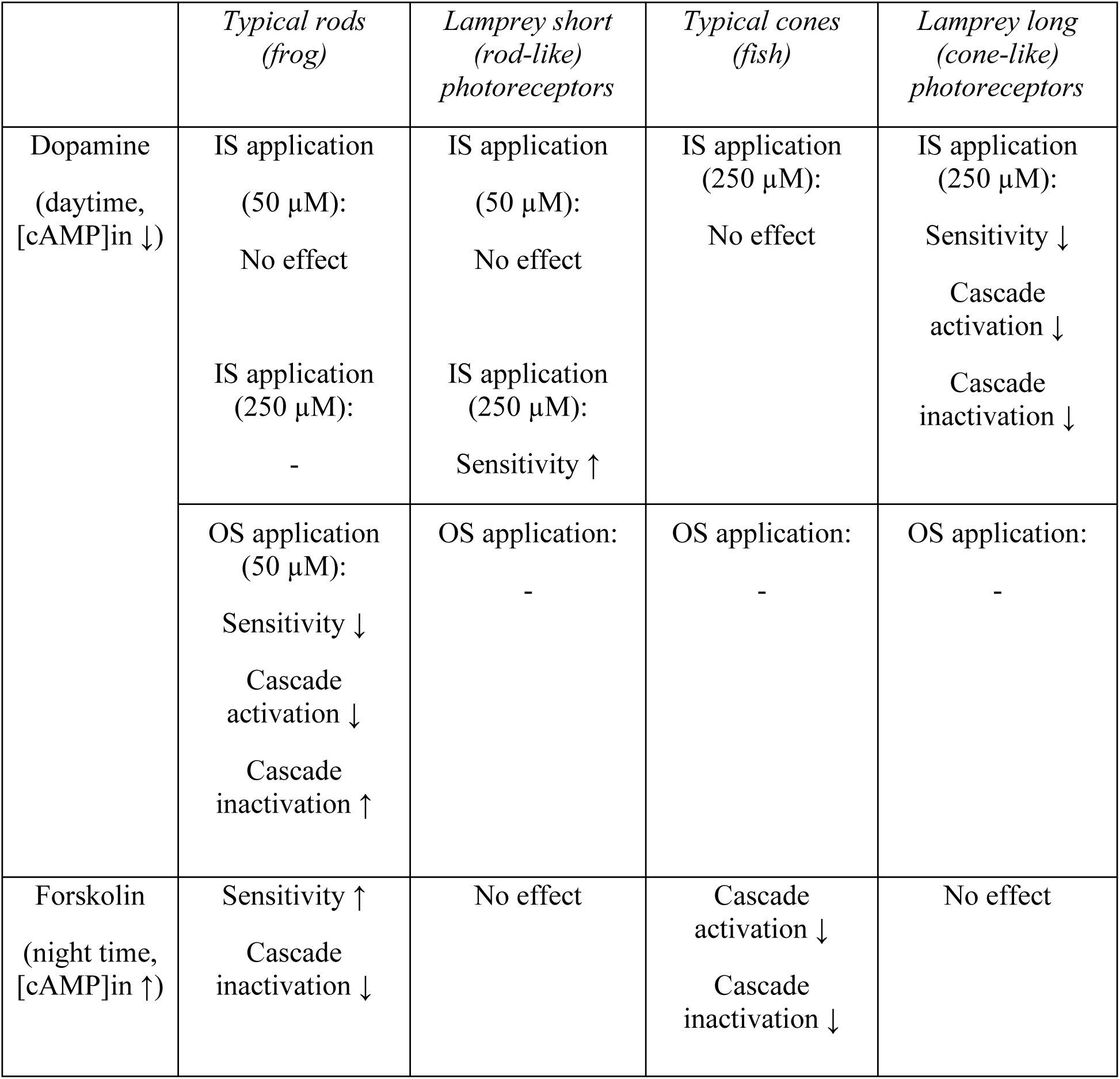
Summary of the effects of dopamine and forskolin on phototransduction in the photoreceptors of different vertebrates based on single-cell recordings. Abbreviations: OS – outer segment; IS – inner segment; dash – no data available. The data are from the present study and from previously published studies on frog rods (Astakhova et al., 2012; Nikolaeva et al., 2019) and fish cones (Sitnikova et al., 2021). Forskolin acts in the same manner, regardless of whether it is applied to the outer or inner segment (see Astakhova et al., 2012).

|  | *Typical rods*<br>*(frog)* | *Lamprey short*<br>*(rod-like)*<br>*photoreceptors* | *Typical cones*<br>*(fish)* | *Lamprey long*<br>*(cone-like)*<br>*photoreceptors* |
| --- | --- | --- | --- | --- |
| Dopamine<br>(daytime,<br>[cAMP]<sub>in</sub> ↓) | IS application<br>(50 μM):<br>No effect | IS application<br>(50 μM):<br>No effect | IS application<br>(250 μM):<br>No effect | IS application<br>(250 μM):<br>Sensitivity ↓<br>Cascade<br>activation ↓<br>Cascade<br>inactivation ↓ |
|  | IS application<br>(250 μM):<br>- | IS application<br>(250 μM):<br>Sensitivity ↑ |  |  |
|  | OS application<br>(50 μM):<br>Sensitivity ↓<br>Cascade<br>activation ↓<br>Cascade<br>inactivation ↑ | OS application:<br>- | OS application:<br>- | OS application:<br>- |
| Forskolin<br>(night time,<br>[cAMP]<sub>in</sub> ↑) | Sensitivity ↑<br>Cascade<br>inactivation ↓ | No effect | Cascade<br>activation ↓<br>Cascade<br>inactivation ↓ | No effect |

From an evolutionary perspective, lampreys are an incredibly interesting study object for retinal processes, as they are the most primitive vertebrates with a complex, multilayered retinal structure. Although their photoreceptors can be functionally categorized as rods and cones (Asteriti et al., 2015; Morshedian and Fain, 2015), they retain a number of intermediate archaic morphological features (Öhman, 1971; Dickson and Graves, 1979) and biochemical properties of phototransduction cascade proteins (Muradov et al., 2008; Govardovskii et al., 2020). Dopamine also acts as a regulatory neurotransmitter in the lamprey retina, as suggested by the presence of amacrine-producing cells (Dacey, 1990) and numerous cells expressing dopamine receptors (present study).

The regulation of the phototransduction cascade by dopamine in lampreys differs significantly from that in gnathostomes: (1) Dopamine affects the response kinetics and photoreceptor sensitivity during the day by acting on the inner segment; (2) Its effects are opposite to those on gnathostomes’ rods; (3) An increase in the intracellular concentration of cAMP (presumably corresponding to night-time) has no effect on photoresponses. The slight increase in rod sensitivity and the decrease in the adaptive potential of cones when exposed to dopamine under daylight conditions (Witkovsky, 2004) may indicate that the visual system shifts into a resource-conserving mode. This mechanism may be related to lampreys’ specific activity cycle during seasonal migrations, when they burrow into the ground and minimize activity during the day (Hardisty, 2013). Whether a similar regulatory mechanism is characteristic of actively hunting sea creatures remains unclear.

The absence of modulation of the phototransduction cascade at high levels of cAMP at night may indicate different fundamental mechanisms of dopamine regulation or in cyclostomes. Indeed, during their upstream migration to rivers, lampreys stop feeding (Hardisty, 2013) and the functionality of many signalling systems in their body cells gradually deteriorates (Savina et al., 2011). To test this theory, we conducted experiments involving the application of forskolin to lampreys at various stages of the migration cycle, from the initial entry into the river in November– December to the spawning readiness stage in April–May and beyond. Forskolin had no effect on lamprey receptor photoresponses at any stage, which suggests that insensitivity to elevated cAMP levels is a fundamental property of their phototransduction cascade. However, this does not exclude the possibility of dopamine acting by reducing cAMP levels via D1-like receptors. The role of dopamine in regulating cyclostome photoreceptors is clearly distinct from that in other vertebrates and merits further research.

## Acknowledgments

The authors express their gratitude to Natalia Drozdova and Dr. Tatiana Ivanova for help with obtaining and keeping lampreys, to Dr. Anatolii Nikiforov for advice on seasonal changes in lamprey physiology, and to Andrei Zherder for the development of the program in Python for approximating photoresponses with mathematical functions.

## Authors’ contributions

D. A. Nikolaeva: performing electrophysiological experiments, data processing, writing–original draft; A. Y. Rotov: performing electrophysiological experiments, preparation of material for immunohistochemistry, methodology, data processing, writing–original draft; I. Y. Morina: immunohistochemical study and Western blotting, manuscript editing; M. L. Firsov: conceptualization, methodology, manuscript editing; I. V. Romanova: conceptualization, immunohistochemical study and Western blotting, manuscript editing; L. A. Astakhova: electrophysiological experiments, data processing, writing–original draft, conceptualization, methodology, project administration.

