## Supplementary material for "Unusual dopamine-mediated regulation of the phototransduction in lamprey compared to jawed vertebrates": Supplemetal figures S1-S8

**Supplemental materials**

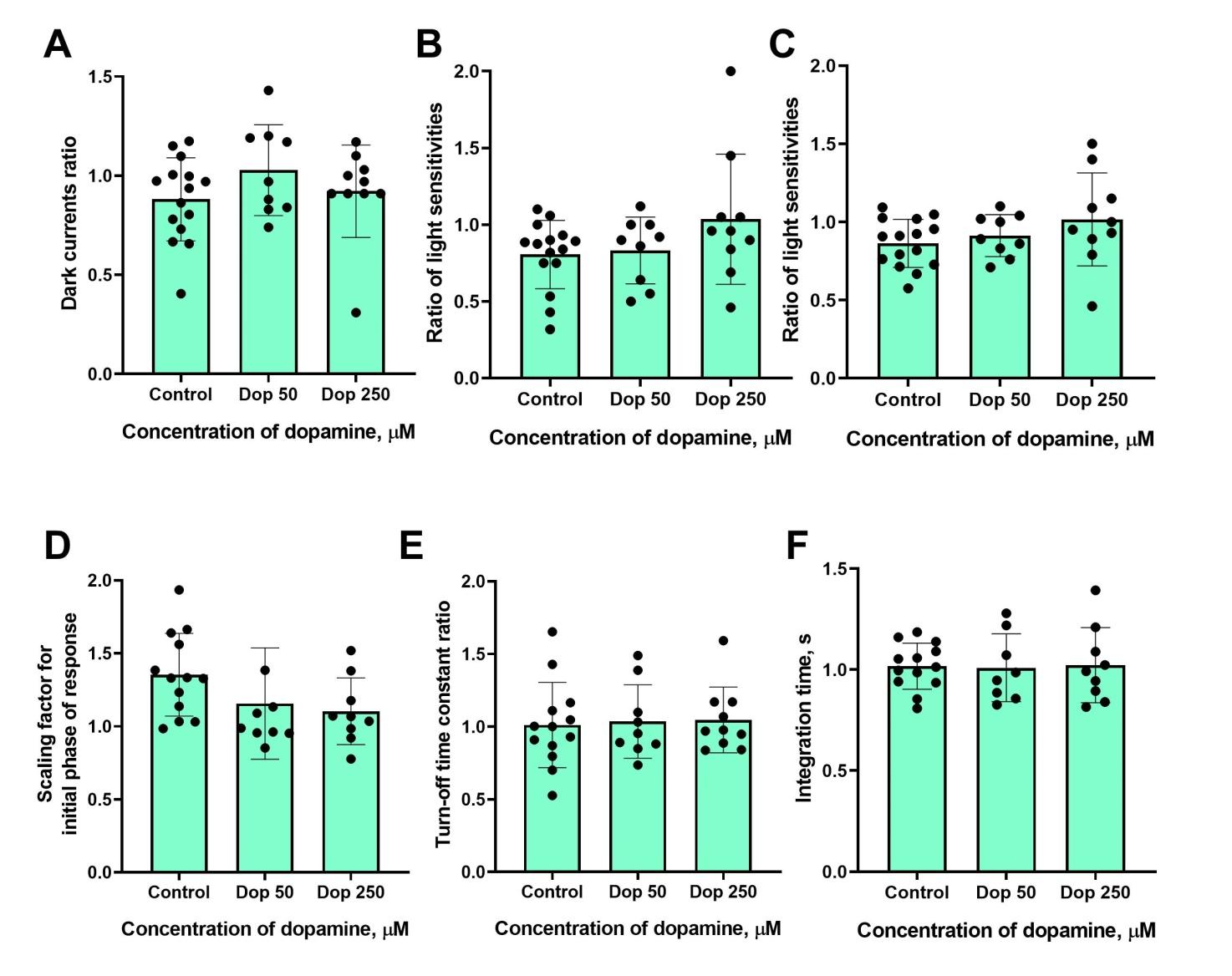

**Figure S1**. Effects of 50 and 250 μM dopamine on the dark current, light sensitivity and photoresponse kinetics of lamprey *short* photoreceptors **after approximately 20 minutes' exposure (first time point in dopamine)**. Comparison of the dark current (A) light sensitivity to quarter-saturating (B) and half-saturating (C) flashes, the scaling coefficient for the rising phase (D), the response recovery time constant (E) and the integration time (F) for responses recorded in normal Ringer's solution and after a 30-minute exposure to dopamine. No statistically significant differences were observed using one-way ANOVA and the post hoc Dunnett test.

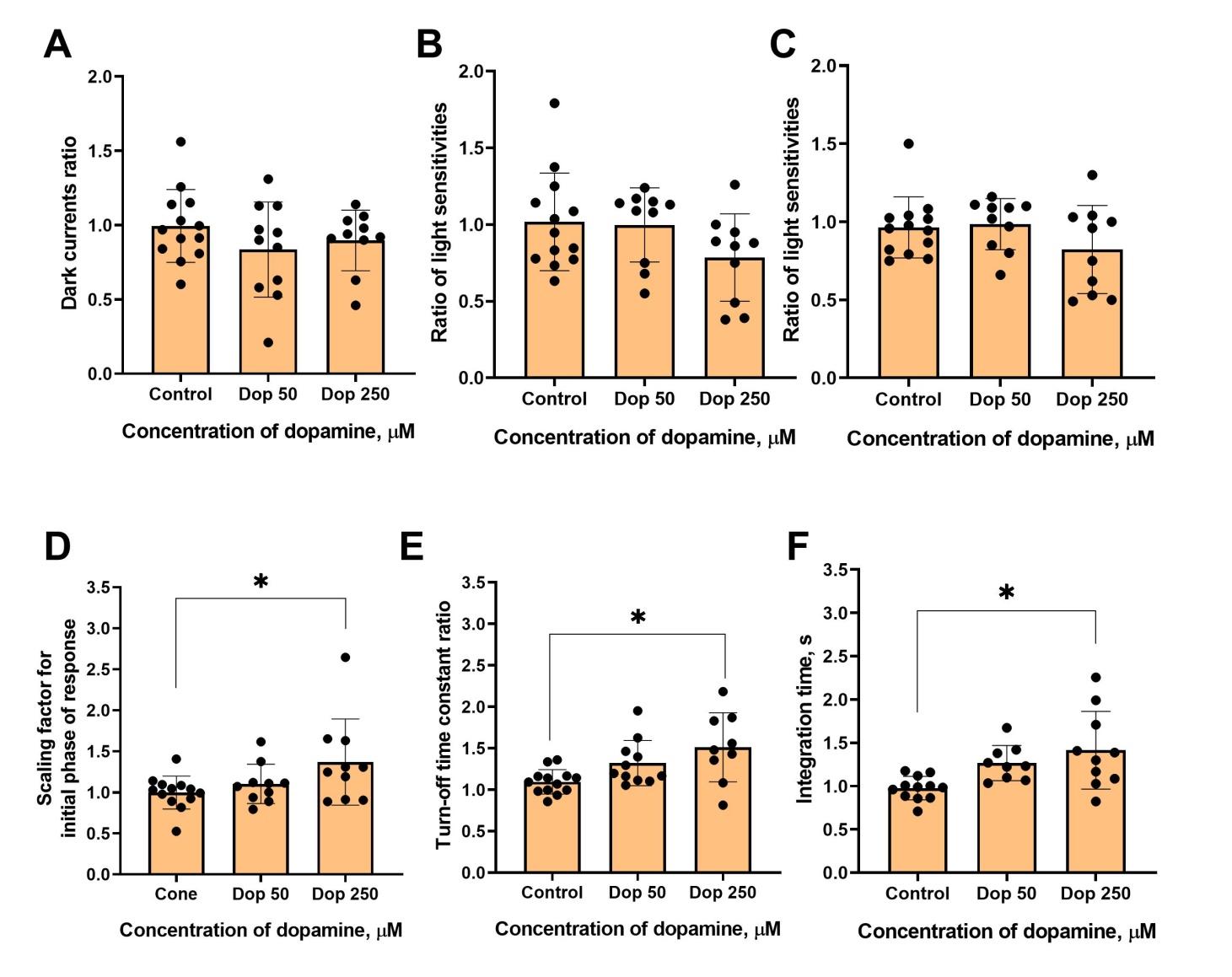

**Figure S2**. Effects of 50 and 250 μM dopamine on the dark current, light sensitivity and photoresponse kinetics of lamprey *long photoreceptors* **after** **approximately 20 minutes' exposure (first time point in dopamine)**. Comparisons of dark current (A) and light sensitivity to near quarter- (B) and half-saturating flashes (C), for responses recorded in normal Ringer's solution and after a 20-minute exposure to dopamine, showed no statistically significant differences (one-way ANOVA and post hoc Dunnett's test). Scaling coefficient for the rising phase (D), response recovery rates (E), and integration time (F) significantly increased for cells incubated in a solution containing 250 µM dopamine compared to the control group (p < 0.05, Dunnett's test).

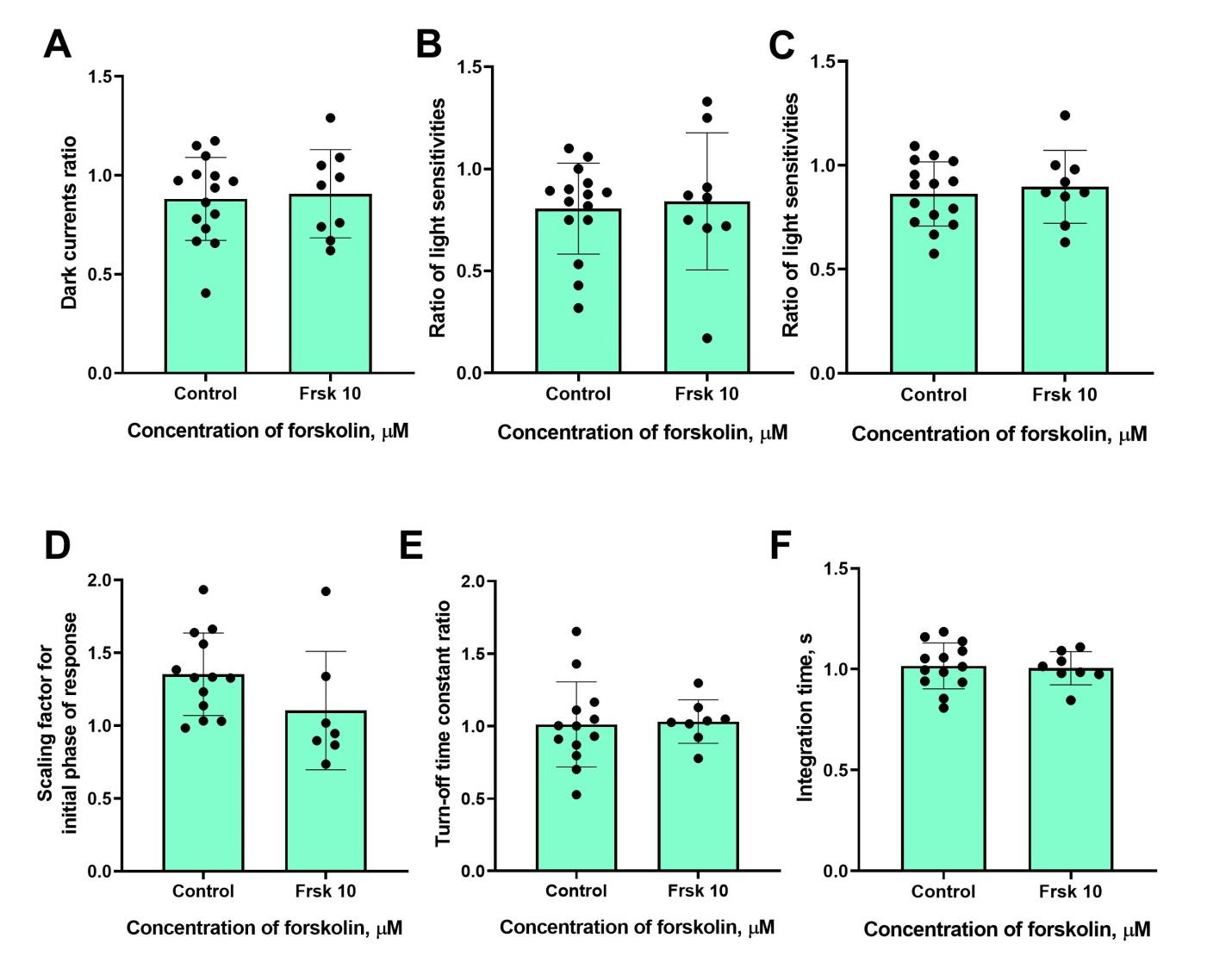

**Figure S3**. The effects of 10 μM forskolin on the dark current, light sensitivity and photoresponse kinetics of lamprey *short photoreceptors* **after approximately 20 minutes' exposure (first time point in forskolin)**. The graph shows a comparison of the dark current (A), light sensitivity to near quarter-saturating (B) and half-saturating (C) flashes, the scaling coefficient for the rising phase (D), response recovery rates (E) and integration time (F) for responses recorded in normal Ringer's solution and after 20 minutes of forskolin exposure. No statistically significant differences were observed using an unpaired t-test.

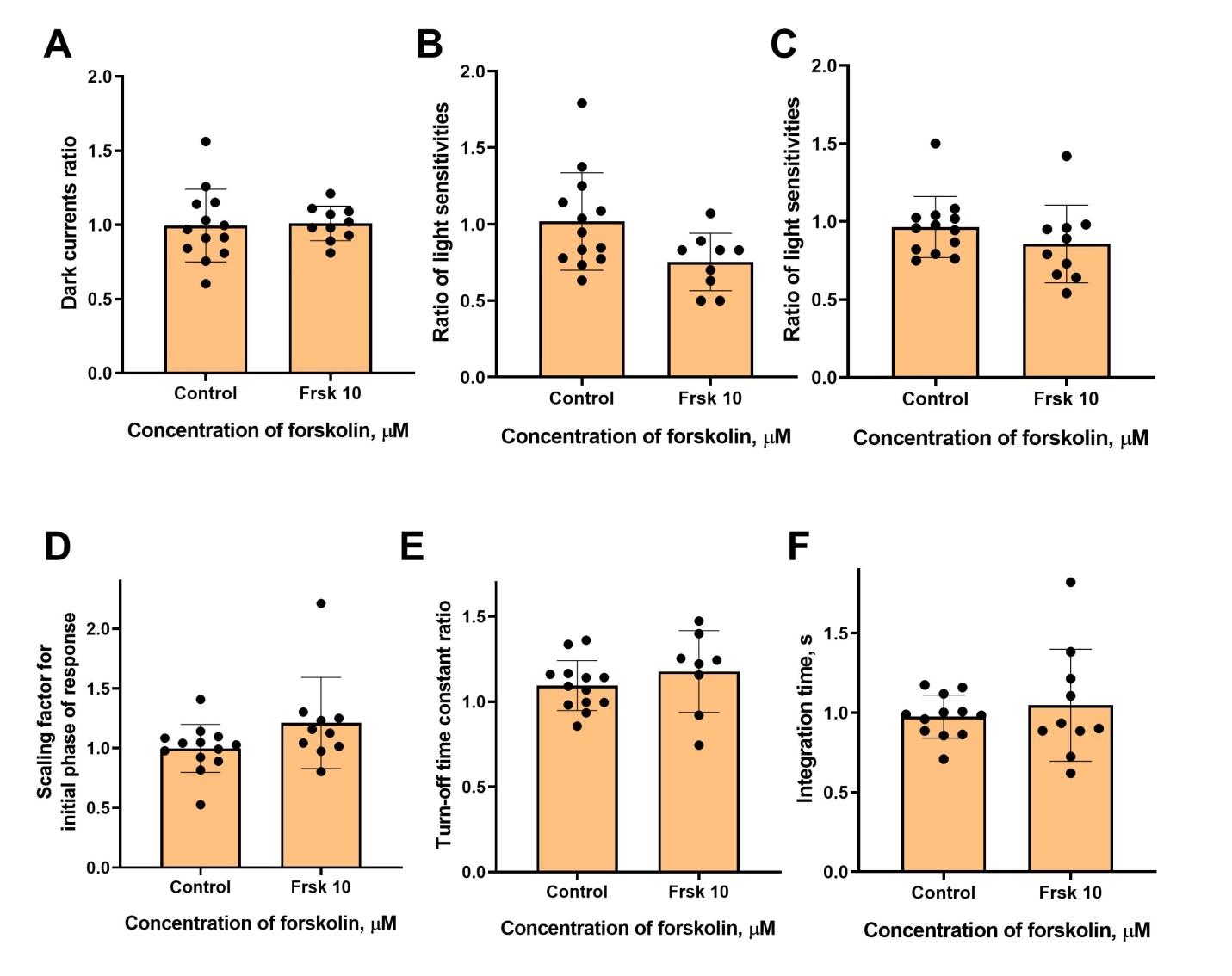

**Figure S4**. The effects of 10 μM forskolin on the dark current, light sensitivity and photoresponse kinetics of lamprey *long photoreceptors* **after approximately 20 minutes' exposure (first time point in forskolin)**. This figure shows a comparison of the dark current (A), light sensitivity to near quarter- (B) and half-saturating flashes (C), the scaling coefficient for the rising phase (D), response recovery rates (E) and integration time (F) for responses recorded in normal Ringer's solution and after 20 minutes of forskolin exposure. No statistically significant differences were observed using an unpaired t-test.

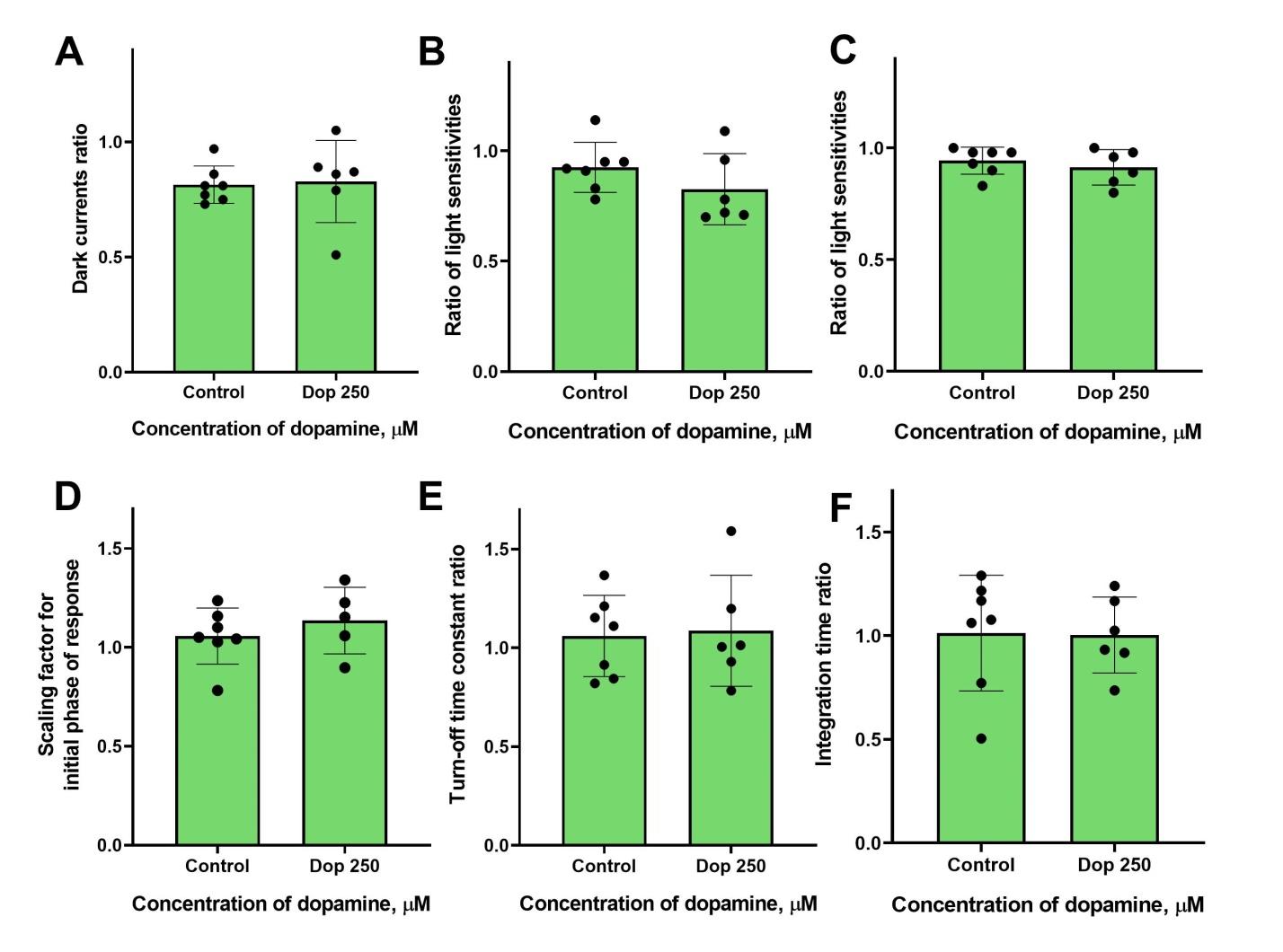

**Figure S5**. Absence of effect of 250 μM dopamine on green-sensitive cones of the fish *Carassius gibelio*: no changes observed after 20 minutes of dopamine exposure in dark current (A) and light sensitivity to near quarter- (B) and half-saturating flashes (C). A comparison of several response kinetic parameters : a scaling coefficient for the rising phase (G), the response recovery rates (H), and the integration time (I) for responses recorded in normal Ringer's solution and after 20 minutes of dopamine exposure. No statistically significant differences were observed using an unpaired t-test.

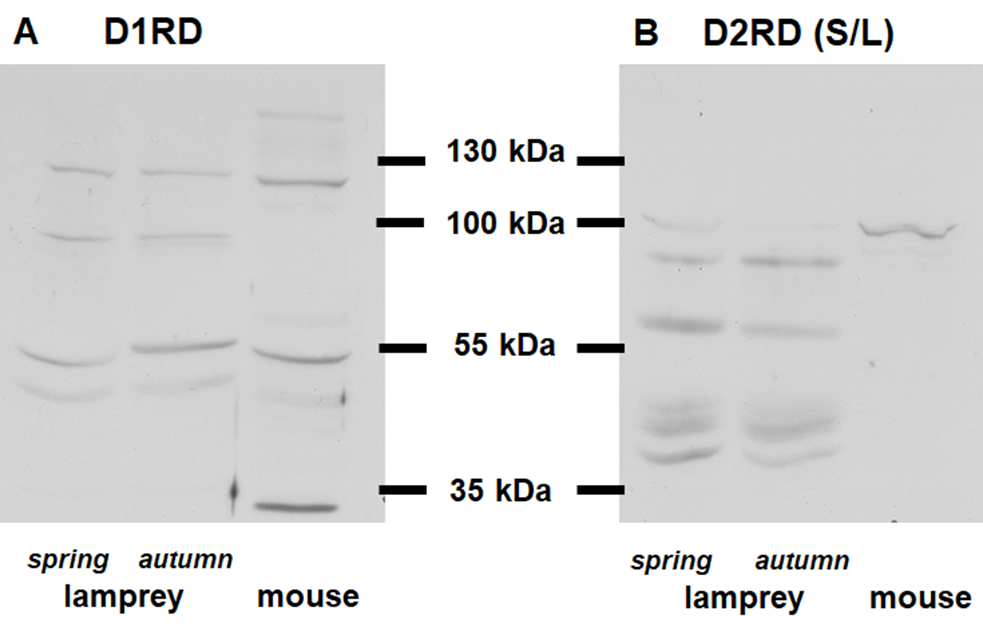

**Figure S6**. Western blotting results demonstrate the presence in the lamprey retina (A) D1RD-immunopositive bands in regions around 55kDa and between 100 and 130kDa and (B) D2RD-immunopositive bands in the 100kDa region and additional bands in the 55kDa region. The mouse retina was used as a positive control. Lamprey retinas for Western blotting were obtained in both spring and autumn to detect potential seasonal variations; however, no differences were observed between these two time points.

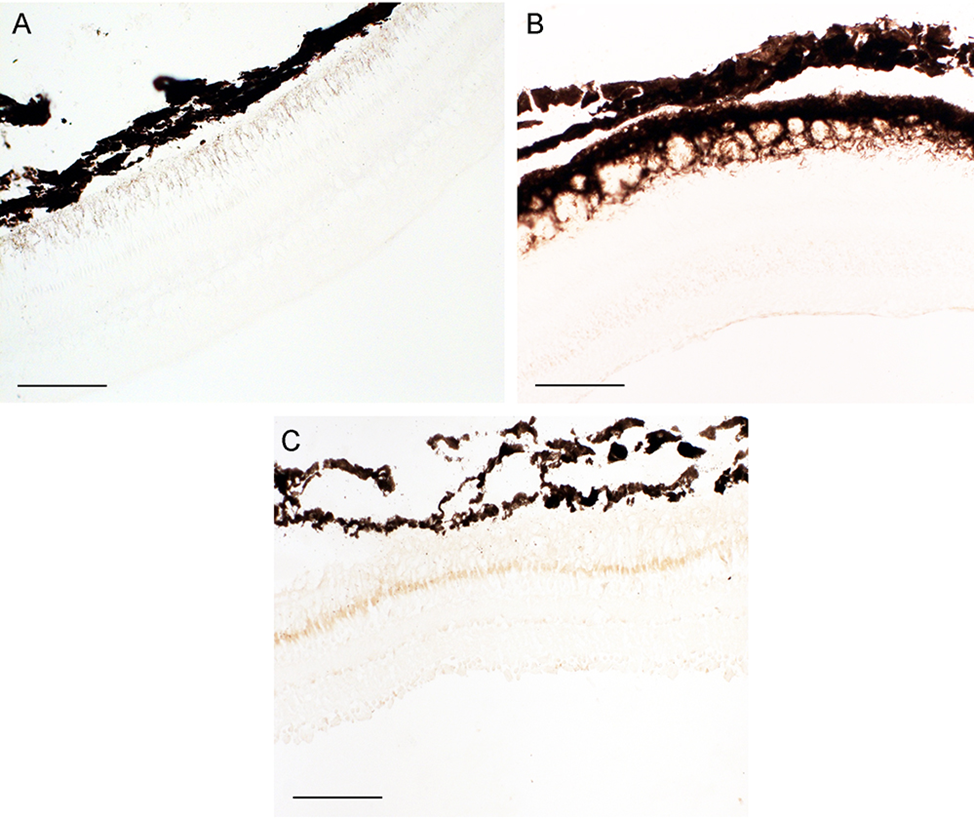
**Figure S7**. Negative controls for immunohistochemical reactions (i.e. reactions without primary antibodies) in the retinas of lampreys (A), carasius (B) and frogs (C). Scale bars: 100 µm.

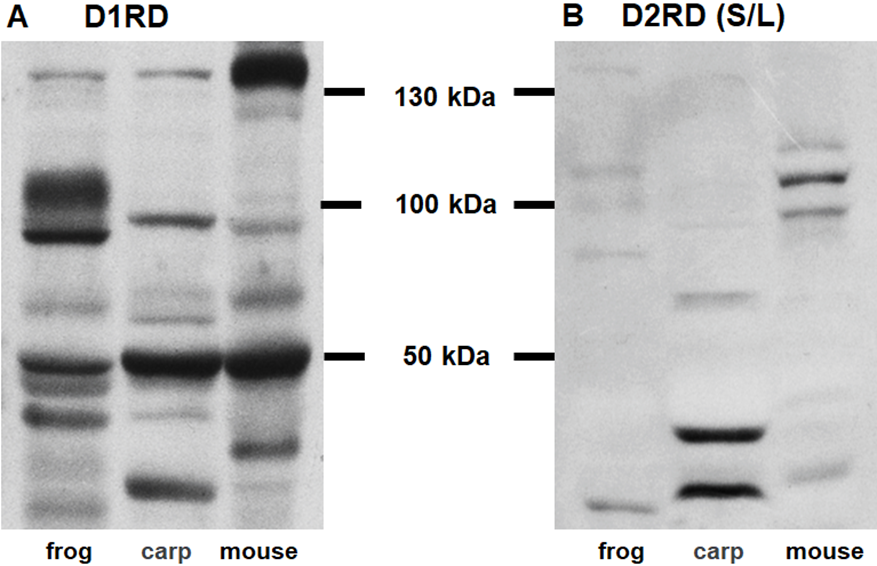

**Figure S8.** Figure S8: Western blotting results demonstrate the presence of immunopositive bands in the retinas of frogs and carasius: (A) D1RD-immunopositive bands in regions around 50, 100 and 130kDa; (B) D2RD-immunopositive bands in an area near 100kDa. The mouse retina was used as a positive control.
